# Frequency-Dependent Modulation of the Prefrontal Cortex by Low-Intensity Focused Ultrasound: Impact on Mesolimbic Dopamine Signaling

**DOI:** 10.1101/2025.10.26.684693

**Authors:** Greatness O. Olaitan, Akhabue K. Okojie, Wendy J. Lynch, B. Jill Venton

## Abstract

Synchronized neural oscillations are fundamental for cognitive function and orchestrate inhibitory and excitatory neurotransmission and downstream signaling. We hypothesized that low-intensity focused ultrasound (LIFU) with frequency parameters mimicking oscillatory patterns would enable targeted neuromodulation. Building on prior findings of the ability of LIFU to modulate neurotransmission, we investigated the effects of frequency-modulated LIFU applied to the prelimbic cortex (PLC) on dopamine release in the nucleus accumbens (NAcc) core as well as neuronal and astrocytic activity. After three 80-second LIFU stimulations spaced 30 minutes apart, we found that LIFU excited or inhibited NAcc dopamine release for up to 90 minutes, and these effects were dependent on oscillation frequency and sex. In male rats, theta (8 Hz)-coupled beta (16 Hz) LIFU reduced both dopamine release by 58% and the level of the astrocytic marker GFAP by 50%. Similar decreases in dopamine density were observed in females. Delta (2 Hz)-coupled beta (16 Hz) stimulation produced similar inhibitory effects. Conversely, theta (5 Hz)-coupled gamma (50 Hz) LIFU increased dopamine release by 28% in males. However, similar excitation levels were observed only in females with an increased gamma frequency of 70 Hz (coupled with a theta frequency of 7 Hz). Histological analysis revealed no cell death, but both 8:16 Hz and 5:50 Hz LIFU elevated neuronal activation (cFOS) in the PLC in males, while 5:50 Hz also upregulated GFAP by 30%, suggesting astrocytic involvement. Thus, LIFU stimulation of the PLC can be frequency-tuned to selectively excite or inhibit dopaminergic signaling in the NAcc, suggesting a novel approach for manipulating neurotransmission.

## Introduction

The electrical activity of the brain exhibits significant coherence, attributed to synchronized neural oscillations across various regions.^1^ These rhythmic fluctuations vary in frequency and demonstrate functional specificity.^2^ EEG research has identified distinct neural oscillatory patterns across specific frequency bands, delta (1–4 Hz), theta (4–8 Hz), alpha (8–12 Hz), beta (13–30 Hz), and gamma (30–150 Hz), that organize neural activity both locally and across brain regions.^2^ Changes in amplitude during large-scale oscillatory activity reflect local synchronization among neural ensembles.^3^ In addition to local synchronization, cross-frequency coupling between neurons in different brain regions facilitates higher-order cognitive functions such as information transfer, perception, motor control, and memory.^4,5^ Disruptions in neuronal synchronization can lead to neural isolation, reduced signal propagation, and cognitive dysfunction.

Neural coupling plays a critical role in regulating various processes and neurotransmitter systems.^6,7^ Among neurotransmitter systems, the mesolimbic dopaminergic pathway is particularly significant due to its role in reward processing and reinforcement learning.^8,9^ Originating in the ventral tegmental area (VTA) and projecting to the nucleus accumbens (NAc) and other limbic structures, this pathway is modulated by the prefrontal cortex (PFC).^10^ The prelimbic cortex (PLC), a subregion of the medial PFC, sends dense glutamatergic projections to the NAc core (NAcc) and modulates dopaminergic signaling to influence reward processing, reinforcement learning, and motivated behavior. The infralimbic cortex (ILC) targets the NAc shell (NAcs), coordinating behavioral motivation and reward-related decision-making.

The oscillatory dynamics of the PFC play a critical role in shaping network activity and downstream signaling, with distinct rhythms contributing to different behavioral states.^11^ During rest, the PFC predominantly exhibits delta oscillations, which reflect baseline activity and large-scale coordination.^12,13^ In contrast, theta activity emerges during active cognitive states, facilitating connectivity between the hippocampus and PFC to support memory integration and decision-making.^14–16^ Beta and gamma oscillations are associated with attentional processes, but they play distinct roles.^17,18^ Beta oscillations exert inhibitory control by stabilizing working memory during delays, clearing irrelevant information after trials, and suppressing memory retrieval during cognitive stopping.^19,20^ Gamma rhythms enhance neuronal synchronization and rapid information processing.^21^ Importantly, cross-frequency coupling between these rhythms further refines PFC function. For example, theta‒gamma coupling enhances synchronization and information transfer.^15,22,23^ These oscillatory dynamics not only shape PFC network activity but also influence signaling in projection regions, such as the NAcc, thereby regulating mesolimbic dopamine pathways.^6,24^ Given that beta oscillations are robustly linked to inhibitory cognitive control and the suppression of irrelevant neural activity,^25^ we reasoned that mimicking theta-beta coupling would engage this inhibitory circuitry, leading to a downstream reduction in dopamine release. Conversely, because theta-gamma coupling is critical for enhancing synaptic potentiation^26^ and information transfer^15,22,23^, we predicted that applying a theta-gamma pattern would amplify excitatory signaling within the PLC, thereby increasing dopamine release in the NAcc.

Building on these predictions derived from oscillatory dynamics, we turned to low-intensity focused ultrasound (LIFU) as a non-invasive method to mimic cross-frequency coupling patterns and probe their causal impact on downstream signaling. We hypothesized that mimicking distinct neural oscillatory patterns via low-intensity focused ultrasound (LIFU) could modulate dopamine signaling in this projection-defined PLC to the NAcc circuit. Previous work from our group demonstrated that brief (2-min) targeted LIFU stimulation of the PLC can induce circuit specific, intensity-dependent inhibition of dopamine release in the NAcc.^27^ However, most LIFU studies have relied on parameters selected based on observed effects rather than testing hypotheses derived from known electrophysiological principles, such as oscillation-coupling dynamics. As an emerging neuromodulatory technique, LIFU offers the spatial specificity and tunability required to systemically probe the relationship between stimulation parameters and circuit-level responses^28^. In particular, pulse repetition frequency (PRF), a LIFU parameter that corresponds to neural oscillation frequency, remains an active area of study for mechanistically informed modulation.

The primary goal of this study was to investigate whether frequency-modulated LIFU targeting of the PLC could selectively influence dopaminergic signaling through its projection to the NAcc. We systematically varied LIFU PRF parameters based on known cross-frequency coupling patterns observed in PFC circuitry. Primarily, we applied LIFU to the PLC and measured NAcc dopamine levels using fast-scan cyclic voltammetry (FSCV). We additionally evaluated neuronal and astrocytic activity via immunohistochemistry, with the ILC-NAc connection serving as a control pathway to contextualize region-specific effects. We hypothesized that while theta-beta coupling would exert an inhibitory effect on dopamine release, theta-gamma coupling amplification would promote greater neuronal synchronization and increase downstream dopamine release in the mesolimbic circuit. Consistent with these hypotheses, our data revealed a 58% decrease in dopamine release, and reduced astrocytic activity was observed under theta (8 Hz)-coupled beta (16 Hz) stimulation. Moreover, there was a 28% increase in dopamine release and increased astrocytic activity under theta-coupled gamma stimulation. We investigated sex differences in the effects of the selected LIFU parameters on dopamine release, neuronal activity, and astrocytic activity. Theta–beta coupling dynamics comparably suppressed dopamine release in both sexes, whereas sustained dopaminergic excitation in females necessitated an elevated gamma oscillation frequency. These results highlight the frequency- and sex-dependent effects of LIFU-induced modulation of the PLC on circuit-specific modulation of dopaminergic signaling. Understanding this relationship provides a systematic approach to optimizing LIFU neuromodulation parameters for treating neuropsychiatric disorders.

## Methods

### Animal surgery

All animal procedures were conducted in compliance with the guidelines set forth by the Animal Care and Use Committee (ACUC) at the University of Virginia. A total of 37 male and 10 female Sprague Dawley rats (Charles River Laboratories, Wilmington, MA, USA), 8–10 weeks of age and weighing between 280 and 320 g, were used in this study, with five rats randomly assigned to each experimental group (sham, anatomical control, 8:16 Hz, 2:16 Hz, 7:28 Hz and 5:50 Hz groups). The sham group had the transducer placed over the PLC, but no ultrasound was delivered. The anatomical control group received the same LIFU stimulation as the treatment groups, but the transducer was aimed at an off-target region, the corpus callosum. The estrous cycle of female rats was not monitored as part of this study’s protocol. The animals were anesthetized using isoflurane, and their core and rectal temperatures were maintained at 36°C throughout the experiment using an isothermal pad (Delta Phase Pad; Braintree Scientific, Braintree, MA, USA). Respiration and anesthesia depth were monitored hourly to ensure stability.

Each rat was secured in a stereotaxic frame, and precise holes were drilled in the skull to position stimulating electrodes, working electrodes, reference electrodes, and the ultrasound transducer following the Paxinos and Watson atlas.^29^ The carbon-fiber working electrode was inserted into the NAcc (+1.3 mm AP, +2.0 mm ML, −7.1 mm DV), while a bipolar stimulating electrode (Plastics One, Roanoke, VA, USA) was placed in the VTA (−4.7 mm AP, +0.9 mm ML, −8.5 mm DV). An Ag/AgCl reference electrode was positioned contralaterally. To target the PLC (+3.6 mm AP, +0.6 mm ML), a 3.0 mm hole was drilled for transducer placement. For anatomical controls, the transducer was angled at 60 degrees to focus on the corpus callosum. Local anesthetic (bupivacaine) was applied to the exposed skin and muscle tissue during surgery to minimize discomfort.

### FSCV measurement of electrically stimulated dopamine

Carbon fiber microelectrodes were made as previously described (Olaitan *et al*., 2025). With a waveform scanned from −0.4 to 1.3 V and back at 400 V/s at 10 Hz, cyclic voltammograms of dopamine were obtained using a WaveNeuro Two FSCV potentiostat (Pine Research, Durham, NC). HDCV software (UNC at Chapel Hill) was used for data acquisition and analysis. Ag/AgCl wires were used as reference electrodes. Carbon fiber microelectrodes were post-calibrated with 1.0 µM dopamine in PBS following the animal experiments. To electrically stimulate dopamine, a constant biphasic current stimulus (24 pulses, +300 μA/pulse, 2 ms pulse width) at 60 Hz and was delivered to the VTA by a bipolar stimulating electrode (Plastics One, Inc., Roanoke, VA, USA). Dopamine was detected using FSCV. Stimulations were delivered every 5 minutes, and dopamine release in the NAcc was measured for 2 hours per LIFU parameter.

### Ultrasound Stimulation

A 10 MHz miniature case immersion transducer (XMS-310-B; Olympus-ims, Waltham, MA) with a 2 mm element diameter and 6 mm focal length was used. Ultrasonic waveforms were generated as described in a previous study by the Venton group. Before LIFU stimulation, the acoustic pressure field was quantified (Figure S1) in an acoustic test tank filled with deionized, degassed, and filtered water (UMS3, Precision Acoustics Ltd., Dorchester, UK) with a calibrated hydrophone (HNR-0500, Onda Corp., CA, USA). Throughout this study, we used a spatial average intensity of 13 W/cm^2^, previously shown to induce downstream dopamine inhibition in the mesolimbic circuit of rats.^27^ The pulse repetition frequency (PRF), interval (PRI), and pulse interval (PI) used are shown in Table 1. Our preliminary experiments showed excitation lasted around 30 minutes but inhibition is longer lasting (Figure S5). We, therefore, administered three successive LIFU trains, separated by 30 minutes, to assess whether the stimulation produced cumulative or lasting neuromodulatory effects on dopamine release dynamics over an extended period.

**Table 1:** LIFU Parameters Optimized for Mimetic Intrinsic Oscillatory Frequency Coupling.

| *Coupling Parameter* | *Abbreviation* | *PRF*<br>*(Hz)* | *Duty Cycle*<br>*(DC)* | *PRI (ms)* | *PI (ms)* | *Expected*<br>*Impact* |
| --- | --- | --- | --- | --- | --- | --- |
| *Theta-Beta (8:16 Hz)* | **BT** | 8 | 50% | 125 | 62.5 | Inhibition |
| *Delta-Beta (2:16 Hz)* | **BD** | 2 | 12.5% | 500 | 62.5 | Inhibition |
| *Theta-High Beta (7:28 Hz)* | **HBT** | 7 | 25% | 142.86 | 35.7 | Excitation |
| *Theta-Gamma (5:50 Hz)* | **GT** | 5 | 10% | 200 | 20 | Excitation |
| *Theta-High Gamma (7:70 Hz)* | **HGT** | 7 | 10% | 142.86 | 14.29 | Excitation |
| *Gamma Only (50 Hz)* |  | 50 | 100% | 20 | 0 | Excitation |

### Histology and Immunofluorescence Staining

After dopamine measurements (40 minutes after the last LIFU stimulation), rats (n =3 per group) from the sham, 16:8 Hz, and 5:50 Hz LIFU groups were first perfused with 1X phosphate-buffered saline (pH = 7.4; Life Technologies Corporation, NY, USA) and then with 4% (wt./vol) paraformaldehyde (PFA) (Product #: 158127; Sigma‒Aldrich, Germany) in phosphate-buffered saline (PBS), after which they were decapitated, and their brains were harvested and postfixed overnight in 4% (wt./vol) PFA. The brains were subsequently washed in PBS and stored at 4 °C. Forty-micron-thick free-floating sections were obtained using a vibratome (Leica VT1000S). To investigate the possibility of LIFU-induced cell death, H&E staining was performed at the University of Virginia histology core as described in a previous study from the Venton group.^27^ For immunofluorescence staining, three sections from each region—the prefrontal cortex (prelimbic and infralimbic) and the striatum (NA core and shell)—were incubated in blocking buffer consisting of 5% normal donkey serum (Product #: 017-000-121; Jackson ImmunoResearch Laboratories, Inc., USA), 0.6% bovine serum albumin (Product #: 9048-46-8; Sigma‒Aldrich, Germany) and 0.3% Triton X-100 solution (Product #: 93443; Sigma‒Aldrich, Germany) in phosphate-buffered saline (PBS) for 2 h at room temperature. Primary antibodies against NeuN (mouse anti-NeuN, 1:1000; product #: MAB377; Sigma‒Aldrich, Germany), c-Fos (guinea pig anti-c-Fos, 1:1000; product #: 226-308; Synaptic System, Germany), and GFAP (chicken anti-GFAP, 1:1000; product #: AB4674; Abcam, United Kingdom) were prepared in blocking buffer, and the sections were incubated with primary antibodies overnight at 4°C. The secondary antibodies used were donkey anti-mouse Alexa Fluor 488 (Product #: 715-545-150; Jackson ImmunoResearch Laboratories, Inc., USA), donkey anti-chicken Alexa Fluor 594 (Product #: 703-585-155; Jackson ImmunoResearch Laboratories, Inc., USA), and donkey anti-guinea pig Alexa Fluor 647 (Product #: 706-605-148; Jackson ImmunoResearch Laboratories, Inc., USA), which were also prepared in blocking buffer (1% NDS, 0.6% BSA, and 0.3% Triton X-100) at a 1:1000 dilution. The sections were then incubated with secondary antibodies for 2 h at room temperature. Following secondary antibody incubation, the sections were washed 3x for 10 minutes each. Sections were treated with an Autofluorescence eliminator (Product #: 2160; Millipore, USA) according to the manufacturer’s protocol. Sections were then mounted with DAPI Fluoromount-G (SB Cat. No. 0100-20). Images were acquired using a Nikon confocal microscope (Nikon Confocal Microscope A1) with a 40× water immersion objective. Cell (NeuN and c-Fos) counting and colocalization and GFAP expression were performed using ImageJ software (version 1.54).

### Statistics

All the statistical analyses were performed using IBM SPSS Statistics (version 21). The p values were significant at the 95% confidence level (*p* < 0.05) and adjusted using the Sidak method. The values in the plots are reported as the means ± standard errors of the means (SEMs), with *n* representing the number of animals. Mixed-effect analyses were also computed to examine group effects and group versus phase interaction effects of LIFU stimulation on the data from different experimental time points.

## Results

### Theta-Beta Frequency (LIFU) of the PLC Inhibits Dopamine Release in its Mesolimbic Projection Target

As a first step toward testing our hypothesis that prefrontal oscillatory dynamics can gate mesolimbic dopamine signaling, we delivered LIFU to the PLC of males via a mechanism tuned to theta–beta coupled rhythms (8:16 Hz). We reasoned that mimicking endogenous theta–beta patterns would entrain PLC activity, leveraging beta’s well-known role in inhibitory control and thereby dampening dopamine release in the NAcc. The experimental arrangement—with the LIFU transducer aimed at the PLC and recording electrode in the NAcc—is described in detail in the methods section (Figure 1A). Three 80-second LIFU trains (8:16 Hz), each separated by 30 minutes, were applied between electrically stimulated dopamine release.

**Figure 1:**
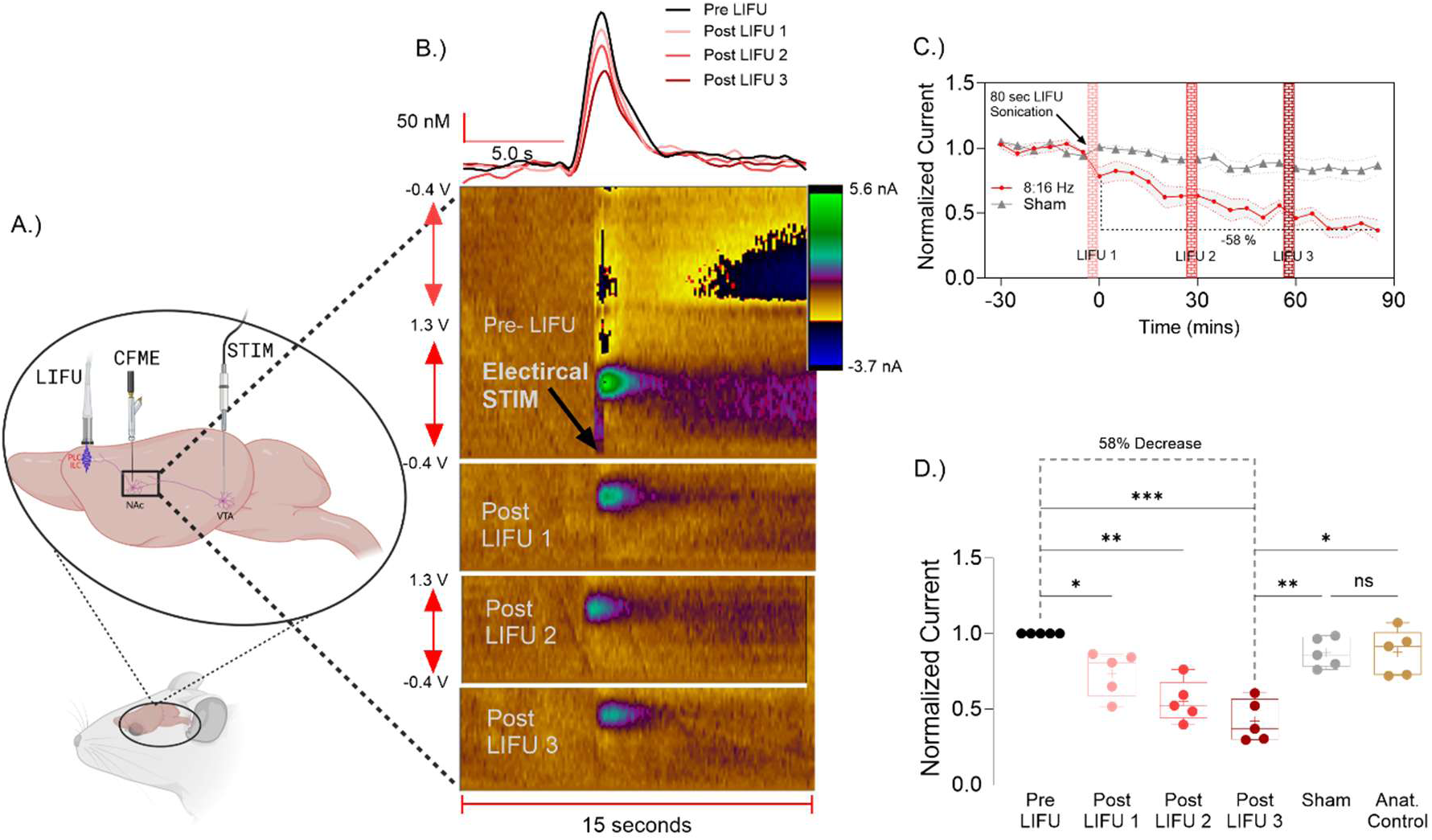
Theta-Beta Frequency LIFU of the PLC Inhibits Dopamine Release in its Mesolimbic Projection Target. **(A)** Schematic of the experimental setup showing the LIFU transducer aimed at the prelimbic cortex (PLC), the carbon-fiber recording electrode in the nucleus accumbens (NAcc), and the stimulating electrode in the medial forebrain bundle. Three 80-s theta–beta-coupled LIFU trains (8:16 Hz) were delivered at 30 min intervals. **(B)** Representative FSCV color plots (bottom) and corresponding concentration-time traces (top) for electrically evoked dopamine release in the NAcc before (pre-LIFU) and after each LIFU train (LIFU 1–3). The color plots demonstrate changes in dopamine release over 15 seconds, with a progressive reduction in signal intensity after LIFU treatment. Dopamine release decreased from 191 ± 10 nM at baseline to 97 ± 10 nM after the third LIFU session. **(C)** Time course of the normalized dopamine-related current (normalized DA; mean ± SEM) sampled every 5 min before (Phase 1) and during three successive 5 min LIFU sessions (Phases 2–4) in the 16-8 Hz (n = 6), Anat (n = 5) and Sham (n = 5) groups. A repeated-measures ANOVA revealed significant main effects of Group (F_(2,12)_ = 11.15, p = .002) and Phase (p < .001) and a Group × Phase interaction (F_(6,36)_ = 7.04, p = .009). Collapsed across phases, the 16-8 Hz group (Δ = 0.677 ± 0.047) produced lower normalized dopamine currents than did the Anatomical (Δ = –0.289 ± 0.066, p = .003) and Sham (Δ = –0.246 ± 0.066, p = .009) groups, whereas the Anatomical and Sham groups did not differ. Phase-specific univariate tests revealed emerging group differences at Phase 2 (F = 6.68, p = .011) and maximal divergence at Phase 4 (F = 13.41, p < .001), with 16-8 Hz significantly lower than that of both controls (Δ = –0.528 to –0.423, p ≤ .006). **(D)** Boxplots of normalized dopamine levels across groups. Multivariate Pillai’s trace confirmed significant within-subject phase effects for 16-8 Hz (F_(3,10)_ = 24.48, p < .001) but not for Anatomical or Sham, and within-subject contrasts demonstrated a pronounced decrease in 16-8 Hz from Phase 1 to Phase 4 (Δ = 0.581 ± 0.076, p < .001), while the control groups remained stable (p > .05). In Phase 4, the 16–8 Hz group differed from the Anat group (Δ = –0.528*, F_(2,12)_ = 13.41, p < .001) and the Sham group (Δ = –0.423*, F_(2,12)_ = 13.41, p = .006), while the Anat versus Sham group was not significantly different (Δ = 0.106, p = .722).

Progressive dopamine suppression was observed across successive post-LIFU phases, with dopamine release decreasing from an average of 191 ± 10 nM to 97 ± 10 nM after the third LIFU application. (Figure 1B). In the 8:16 Hz group, the normalized evoked dopamine concentration decreased by 58% cumulatively over the three sessions, whereas the sham controls maintained stable release profiles (Figure 1C). A repeated-measures ANOVA revealed a significant group × phase interaction, with dopamine in the 8:16 Hz cohort dropping to less than half of baseline by the final phase. Multivariate analyses (Table S1E) confirmed the time-dependent escalation of suppression unique to the LIFU-treated animals, and within-group contrasts underscored sustained inhibition over the entire poststimulation period (Figure 1D, Table S1D). The LIFU group differed significantly from both controls, and the anatomical group did not differ from the no-LIFU control group (Figure 1D), validating both the frequency- and anatomy-specific actions of LIFU. These data demonstrated that theta–beta coupled LIFU effectively and specifically disrupted PLC–NAcc circuit dynamics, producing cumulative neuromodulatory effects that suppressed dopamine release.

### Beta Frequency-Dependent Inhibition of NAcc Dopamine Release by LIFU

Building on the well-characterized relationship between 8:16 Hz LIFU stimulation and dopaminergic inhibition, we expanded our methodological framework to systematically evaluate frequency-dependent inhibitory effects. This was achieved by directly contrasting two distinct coupling paradigms: theta-coupled beta (8:16 Hz) and delta-coupled beta (2:16 Hz) stimulation. Both protocols employed a 10 MHz carrier frequency delivered in 80-second trains, differentiated by pulse repetition intervals to generate their respective oscillatory configurations (Figure 2A).

**Figure 2.**
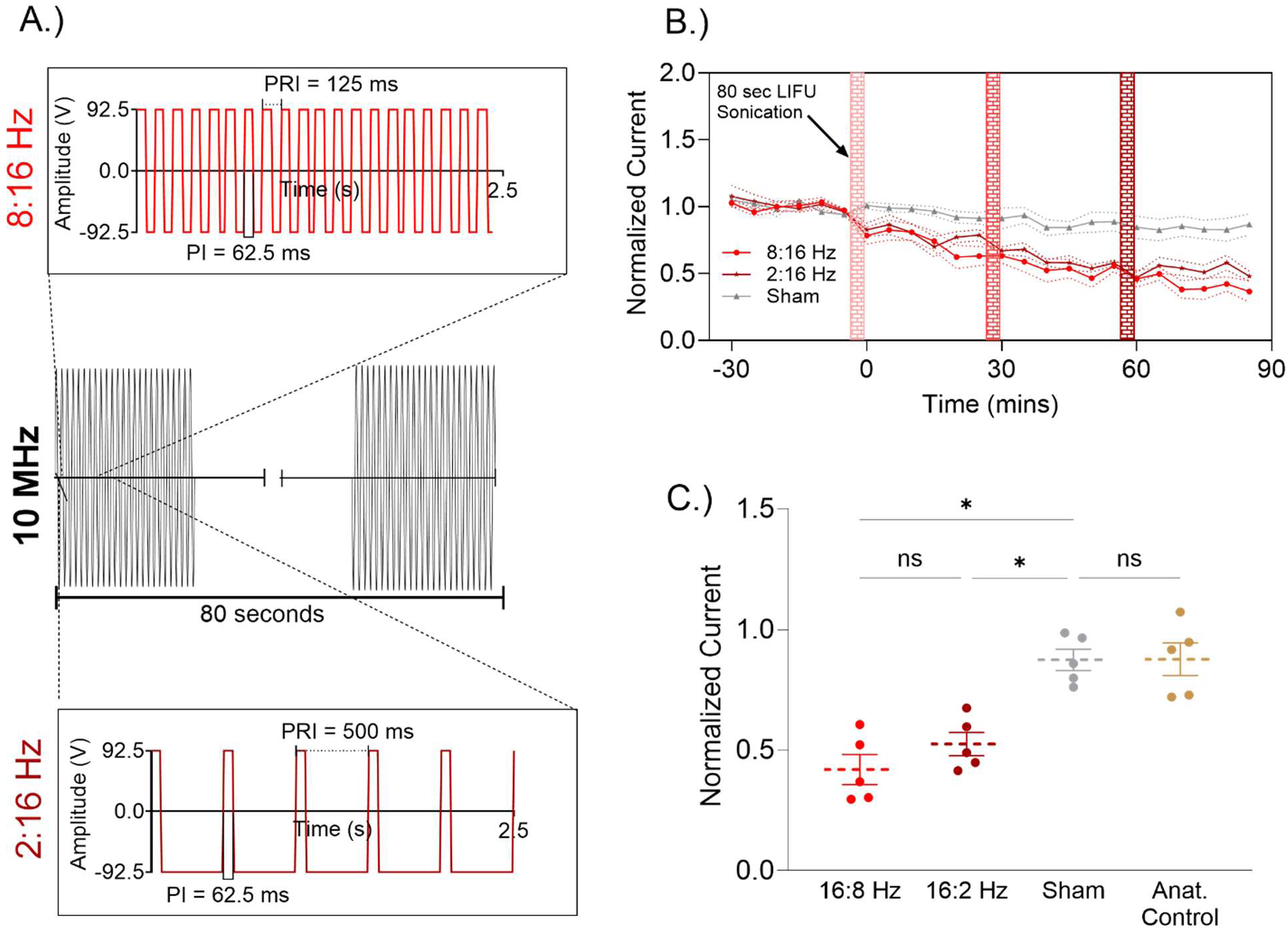
Frequency-dependent inhibition of dopamine release by LIFU. **(A)** Schematic of LIFU pulse parameters: 10 MHz fundamental frequency; pulse-repetition interval (PRI): 125 ms for theta–beta coupling (8:16 Hz) and 500 ms for delta–beta coupling (2:16 Hz); each pulse interval (PI): 62.5 ms; delivered as 80-s trains. **(B)** Time course of the normalized dopamine-related current (mean ± SEM; Anat, n = 5; Sham, n = 5; 2:16 Hz, n = 6; 8:16 Hz, n = 6) sampled every 5 min before (Phase 1) and after (Phase 2–4) LIFU. A mixed-design ANOVA revealed significant main effects of Group and Phase (both p < .001) and a Group × Phase interaction (p < .001). In Phase 1, Anat (M = 0.966 ± 0.039) exceeded BD (Δ = 0.218 ± 0.053, p = .004) and BT (Δ = 0.285 ± 0.053, p < .001) but did not differ from Sham (Δ = 0.044 ± 0.055, p = 0.968). Over Phases 2–4, 2:16 Hz and 8:16 Hz declined to 0.542 and 0.404, respectively, whereas Anat (0.947) and Sham (0.842) remained stable. Multivariate Pillai’s trace confirmed significant phase effects for 2:16 Hz (Trace = 0.785, F_(3,16)_ = 19.512, p < .001) and 8:16 Hz (Trace = 0.878, F_(3,16)_ = 38.473, p < .001) but not for Anat or Sham. In Phase 1 (Pre-LIFU), no differences between groups emerged (all Δ = 0, ns). By Phase 2 (LIFU 1), there was a significant group effect, F_(3,18)_ = 5.199, p = .009, with 8:16 Hz showing a modest but significant reduction in dopamine release relative to Anat (Δ = 0.191 ± 0.063, p = .040) and Sham (Δ = 0.214 ± 0.063, p = .018). In Phase 3 (LIFU 2), the group effect increased (F_(3,18)_ = 8.673, p < .001): 2:16 Hz was lower than that in Anat (Δ = 0.350 ± 0.095, p = .010), and 8:16 Hz was lower than that in Anat (Δ = 0.407 ± 0.095, p = .003) and lower than that in Sham (Δ = 0.316 ± 0.095, p = .022). By Phase 4 (LIFU 3), group differences peaked, F_(3,18)_ = 16.204, p < .001, with Anat vs. 2:16 Hz Δ = 0.405 ± 0.089, p = .001; Anat vs. 8:16 Hz Δ = 0.543 ± 0.089, p < .001; Sham vs. 2:16 Hz Δ = 0.300 ± 0.089, p = .021; and Sham vs. 8:16 Hz Δ = 0.437 ± 0.089, p < .001. **(C)** Boxplots of the mean normalized current across phase 4 trains for each group. Univariate ANOVAs revealed a progressive difference in the normalized dopamine current between the LIFU-treated group and the control group across the four groups. At phase 4 (LIFU 3), the 2:16 Hz (Anat vs. 2:16 Hz, Δ = 0.405 ± 0.089, p = .001; Sham vs. 2:16 Hz, Δ = 0.300 ± 0.089, p = .021) and 8:16 Hz (Anat vs. 8:16 Hz, Δ = 0.543 ± 0.089, p < .001; and Sham vs. 8:16 Hz, Δ = 0.437 ± 0.089, p < .001) groups differed significantly from the Anat and Sham control groups.

Time-course analyses demonstrated that both 8:16 Hz and 2:16 Hz LIFU elicited substantial suppression of the normalized dopamine current relative to that in the sham controls (Figure 2B). While 2:16 Hz stimulation exhibited a modestly greater suppression magnitude than 8:16 Hz stimulation, this difference did not reach statistical significance (Figure 2C). The anatomical control measurements were indistinguishable from the baseline values, confirming the strict spatial specificity of the stimulation effects (Table S2C).

A mixed-design ANOVA incorporating four experimental groups and temporal phases (pre-LIFU + three post-LIFU phases) revealed significant main effects of Group and Group × Phase interaction (Table S2C). Dopamine release was significantly greater in the Anat and sham groups than in the 2:16 Hz and 8:16 Hz groups, while the release in the Anat controls was statistically equivalent to that in the Sham controls. Longitudinal analysis of poststimulation phases demonstrated progressive response attenuation in the 2:16 Hz group, in contrast to the stable trajectories observed in the Anatomical and Sham groups. By Phase 4, both the 2:16 Hz and 8:16 Hz FOPs exhibited significant suppression relative to the Anatomical and Sham conditions (Table S2C). Multivariate analyses (Pillai’s trace) corroborated significant phase-dependent effects exclusively in the 2:16 Hz and 8:16 Hz groups (Table S1E). These findings establish that delta-and theta-coupled beta LIFU paradigms induce robust, frequency-dependent (although nonselective) inhibition of NAcc dopamine release.

### Theta-Gamma LIFU to Upregulate the Release of PLC-Induced NAcc Dopamine

To further elucidate the role of prefrontal oscillatory dynamics in gating mesolimbic dopamine transmission, we applied 80-second trains of LIFU to the PLC, precisely tuned to theta-coupled gamma rhythms (5:50 Hz). Initial stimulation with a one-hour interstimulation interval (ISI) induced transient elevations in dopamine release, which returned to baseline within 30 minutes (Fig. S5A). To enhance temporal stability, we reduced the ISI to 30 minutes and continuously recorded electrically evoked dopamine release in the NAcc.

This revised paradigm yielded sustained potentiation of dopamine signaling. Representative FSCV color plots and concentration-time traces illustrate persistent augmentation of dopamine release across all three LIFU sessions (Fig. 3A). Dopamine concentrations increased from a pre-LIFU baseline of 154 ± 4 nM to 215 ± 4 nM, maintaining an approximate 28% increase throughout the study period, without evidence of synaptic fatigue or depletion commonly associated with repeated stimulation paradigms (Fig. 3A inset). Notably, animals receiving 5:50 HzLIFU exhibited robust and sustained dopamine enhancements across post-LIFU phases (Fig. 3B). Dopamine release increased by 30% in Phase 2, remained elevated at 28% in Phase 3, and persisted at 26.5% in Phase 4, each of which significantly exceeded the responses observed in the anatomical and sham control groups. A mixed design analysis revealed a significant main effect of group on dopamine release (Table S3C), with the 5:50 Hz cohort demonstrating elevated dopamine levels relative to both anatomical and sham controls, between which no significant difference was detected. Phase-specific effects revealed significant within-group modulation in the 5:50 Hz group alone (Table S3D). Multivariate analysis confirmed that the 5:50 Hz cohort had a strong phase effect (Table S3E). Importantly, comparisons against both sham anatomical and off-target stimulation controls demonstrated the anatomical and frequency specificity of this effect. Following the third LIFU train, dopamine released in the 5:50 Hz group remained significantly elevated relative to that in the anatomical controls (Fig. 3C), whereas dopamine released in the anatomical and sham controls was not significantly different. Collectively, these findings demonstrate that theta–gamma–coupled LIFU to the PLC drives temporally extended potentiation of mesolimbic dopamine release. This enhancement persists across repeated stimulations without inducing typical depletion, supporting the hypothesis that frequency-specific neuromodulation can reinforce dopaminergic circuit resilience.

**Figure 3:**
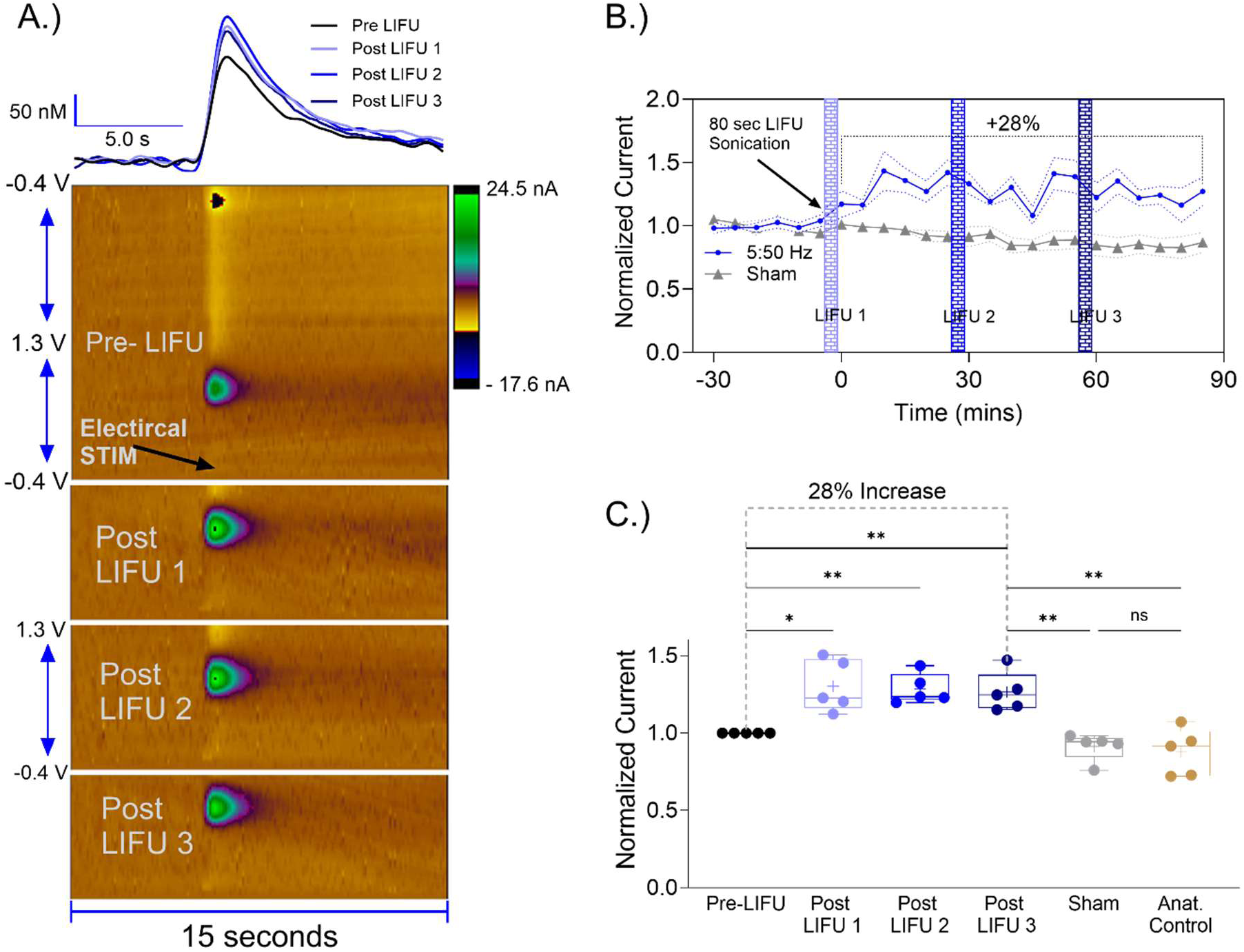
Theta-Gamma LIFU-induced Sustained Dopamine Release Upregulation in the NAcc. **(A)** Representative FSCV color plots (bottom) and corresponding concentration-time traces (top) illustrating electrically evoked dopamine release in the nucleus accumbens before and after three 80-s theta–gamma (5:50 Hz) LIFU trains were delivered to the prelimbic cortex at 30-min intervals. Poststimulation traces reveal a clear and lasting increase in dopamine oxidation currents over the 15-s sampling window. Overall, the dopamine amplitude increased from 154 ± 4 nM pre-LIFU to 215 ± 4 nM following the final stimulation. **(B)** Time course of the normalized dopamine current (mean ± SEM; n = 5 per group) sampled every 5 min before (Phase 1) and after each LIFU train (phases 2–4) for both the 5:50 Hz and anatomical control groups. Dopamine levels increased by an average of ∼28% above baseline over the course of the 90 min post-LIFU stimulation. Mixed model analysis revealed significant main effects of Group (F(2,12) = 12.53, p = 0.001) and Phase (p < 0.001) and a Group × Phase interaction (Pillai’s trace = 0.751, F(3,10) = 10.06, p = 0.002). When collapsed across phases, the 5:50 Hz cohort exhibited greater normalized currents (Δ = 0.677 ± 0.047) than did the anatomical controls (Δ = –0.289 ± 0.066; p = 0.006). Univariate tests confirmed emerging group differences at Phase 2 (F(2,12) = 14.83, p < 0.001) and maximal divergence at Phase 4 (F(2,12) = 8.65, p = 0.005), with the 5:50 Hz group being significantly elevated relative to anatomical controls (Δ = 0.528–0.423, p ≤ 0.006). **(C)** Boxplots of normalized dopamine (NormalizedDA) across experimental conditions. Mixed-design ANOVA indicated a significant group effect (F(2,12) = 12.53, p = 0.001); the theta– gamma (5:50 Hz) group exceeded both the anatomical (p = 0.006) and sham (p = 0.002) controls, which did not differ (p = 0.874). The Group × Phase interaction (Pillai’s trace = 0.751, F(3,10) = 10.06, p = 0.002) reflected consistent within-group potentiation across Phases 2–4 (Phase 2: p < 0.001; Phase 3: p = 0.024; Phase 4: p = 0.024, Sidak-adjusted). The sham and anatomical control groups remained stable over time (p > 0.24). Notably, in Phase 4, 5:50 Hz dopamine release was significantly greater than that in the sham group (p = 0.0054), confirming both the frequency- and anatomy-specific effects of LIFU.

### Theta–Gamma Coupling Drives LIFU-Induced Dopamine Release via Neuronal Synchronization

Building on our demonstration that theta–gamma LIFU potentiates NAcc dopamine, we asked whether this enhancement reflects genuine neuronal synchronization rather than merely a longer pulse interval. To address this, we directly contrasted two coupling paradigms—theta–gamma (5:50 Hz) versus theta–high–beta (7:28 Hz) LIFU—each delivered as 80-s, 10 MHz carrier trains but differing in PRI/PI (5:50 Hz: 200 ms PRI, 20 ms PI; 7:28 Hz: 142.86 ms PRI, 35.71 ms PI; Fig. 4A).

**Figure 4.**
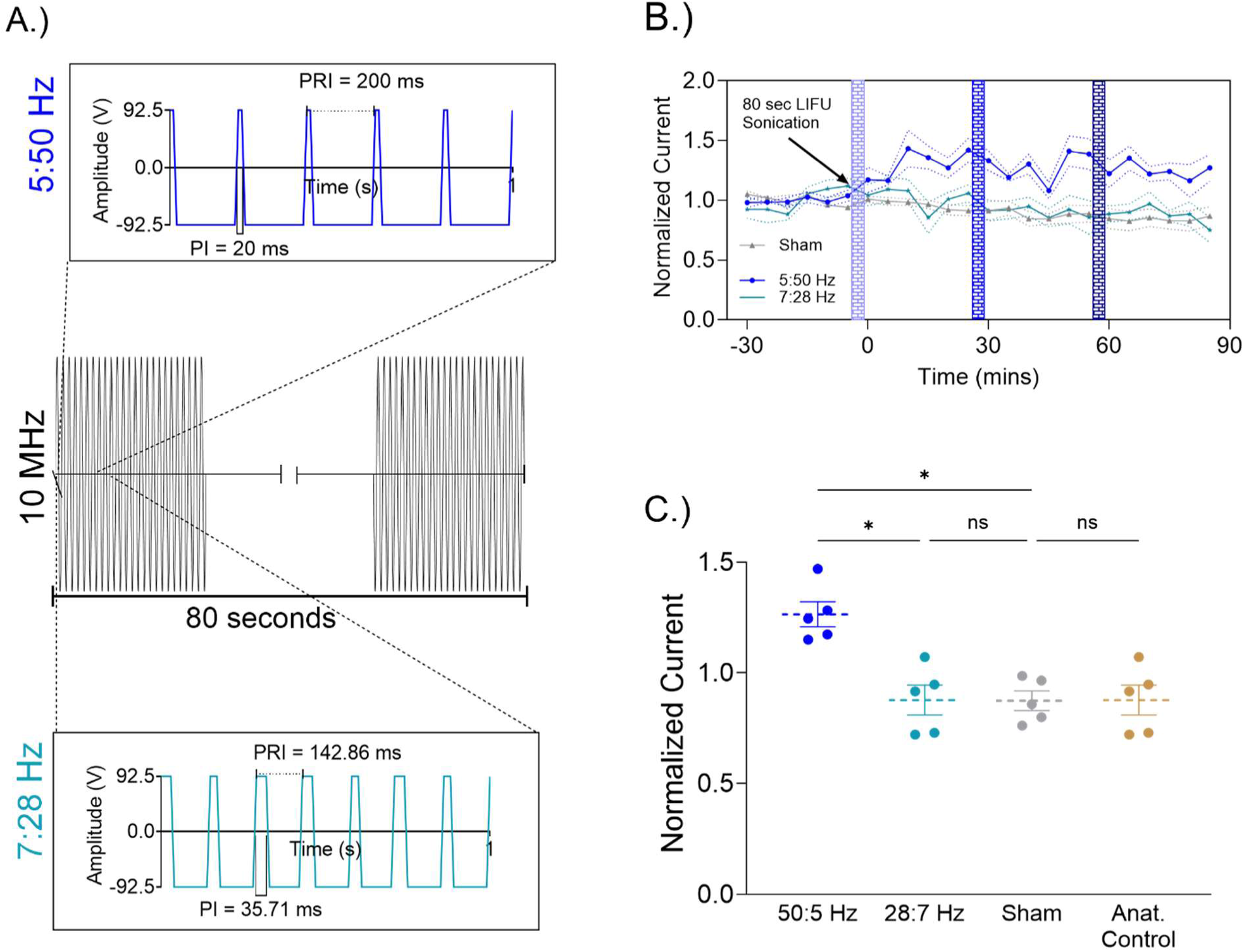
Theta–Gamma Coupling Drives LIFU-Induced Dopamine Release via Neuronal Synchronization. **(A)** Schematic of the two LIFU paradigms, each delivered as an 80-s, 10 MHz carrier train to the prelimbic cortex. Insets show the pulse waveform details. The blue graph shows the 5:50 Hz LIFU waveform (pulse repetition interval (PRI) = 200 ms, pulse interval (PI) = 20 ms, amplitude ± 92.5 V, time axis = 100 ms). The teal line graph shows the 7:28 Hz LIFU waveform (PRI = 142.86 ms, PI = 35.71 ms, amplitude ± 92.5 V (time axis = 100 ms). **(B)** Time course of the normalized dopamine-related current (mean ± SEM; Anat, n = 5; Sham, n = 5; 5:50 Hz, n = 6; 7:28 Hz, n = 6) sampled every 5 min before (Phase 1) and after (Phases 2–4) LIFU. A mixed-design ANOVA revealed significant main effects of Group (F(3,16) = 5.51, p = 0.009) and Phase (p < 0.001) and a Group × Phase interaction (Fs(3,16) = 5.505–4.837, p < 0.05). In Phase 1, Anat (M = 0.966 ± 0.058) did not differ from Sham (Δ = 0.044 ± 0.081, p = 0.996), 7:28 Hz (Δ = 0.019 ± 0.081, p = 1.000) or 5:50 Hz (Δ = –0.247 ± 0.081, p = 0.047). Over Phases 2–4, 5:50 Hz exhibited sustained elevation (Phase 2: M = 1.302; Phase 4: M = 1.265), whereas 7:28 Hz declined progressively to 86% by Phase 4. Anat (0.947) and Sham (0.842) mice remained stable. Multivariate Pillai’s trace confirmed significant phase effects at 5:50 Hz (trace = 0.542, F(3,14) = 5.513, p = 0.010) but not at Anat, 7:28 Hz, or Sham. By Phase 2, a significant group effect (F(3,16) = 5.935, p = 0.006) was driven by 5:50 Hz> Anat (Δ = 0.360 ± 0.099, p = 0.013) and 5:50 Hz> Sham (Δ = 0.338 ± 0.099, p = 0.021). In Phase 3, 5:50 Hz remained elevated compared with that in the Sham group (Δ = 0.400 ± 0.129, p = 0.041). In Phase 4, the 5:50 Hz bandpassed 7:28 Hz (Δ = 0.404 ± 0.126, p = 0.033) and the Sham (Δ = 0.424 ± 0.126, p = 0.024) band. Within-group contrasts confirmed a significant increase from Phase 1 to Phase 2 for 5:50 Hz (Δ = 0.302 ± 0.070, p = 0.003), whereas 7:28 Hz decreased nonsignificantly in Phase 4 (Δ = –0.139 ± 0.089, p = 0.594). **(C)** Scatterplot of individual normalized dopamine currents with group means ± SEMs. A mixed-design ANOVA showed a significant group effect (F(3,16) = 5.51, *p* = 0.009). Post hoc Sidak tests confirmed that 5:50 Hz exceeded the Anatomical frequency (Δ = 0.247*, *p* = 0.047), 7:28 Hz (Δ = 0.266*, *p* = 0.029) and Sham (Δ = 0.290*, *p* = 0.015), whereas the Anatomical, 7:28 Hz and Sham frequencies did not significantly differ (*p* > 0.99). Univariate phase-specific tests demonstrated sustained elevation only in the 5:50 Hz group (Phase 2: F(3,16) = 5.935, *p* = 0.006; Phase 3: F(3,16) = 3.833, *p* = 0.030; Phase 4: F(3,16) = 4.837, *p* = 0.014). Multivariate analysis (Pillai’s trace) confirmed a selective phase effect at 5:50 Hz (F(3,14) = 5.513, *p* = 0.010) but not for the other groups.

Time-course analyses of FSCV recordings revealed that, compared with the sham controls, only the 5:50 HzLIFU protocol drove a marked and sustained increase in the normalized dopamine current, whereas the 7:28 Hz protocol did not affect the baseline current (Fig. 4B). Anatomical and sham controls were likewise tracked with the 7:28 Hz group, confirming the spatial specificity of theta–gamma entrainment (Fig. 4C). A mixed-design ANOVA across four groups (Anat, 5:50 Hz; 7:28 Hz; Sham) and four phases (pre-LIFU + three post-LIFU intervals) revealed a significant main effect of Group and a Group × Phase interaction (Table S4C). Collapsing across phases, the 5:50 Hz cohort exhibited significantly greater dopamine release than both the Anatomical and Sham controls (Table S4D), while the 7:28 Hz cohort did not differ. Phase-specific univariate contrasts confirmed that only the 5:50 Hz paradigm maintained elevated dopamine release throughout all poststimulation phases. Multivariate tests (Pillai’s trace) further substantiated a selective phase effect for the 5:50 Hz group (Table S4E).

To determine whether gamma oscillations alone could boost dopaminergic output, we evaluated continuous 50 Hz (100% duty cycle) PRF LIFU (Fig. S2), which yielded a modest 21% increase in dopamine. Thus, while gamma entrainment enhances NAcc neurotransmission, the addition of theta coupling amplifies and prolongs this effect. Collectively, these findings demonstrate that theta–gamma synchronization underlies the robust upregulation of mesolimbic dopamine by LIFU.

### Sex-specific modulation of dopamine release by LIFU

To evaluate potential sex differences in LIFU responses, we extended our analysis to female subjects across both the 8:16 Hz and 5:50 Hz stimulation paradigms. At 8:16 Hz, males and females exhibited robust and comparable reductions in the normalized dopamine current of approximately 60% following three successive stimulations, with no significant differences observed between the sexes (Fig. 5A). Multivariate analysis confirmed the significant effects of phase progression in both groups, each of which showed a similar pattern of decline in dopamine release over time (Table S5E). No significant difference was found in the magnitude of dopamine inhibition at any individual phase between males and females (Table S5C), a result visually supported by the overlapping box plots in Figure 5 B.

**Figure 5.**
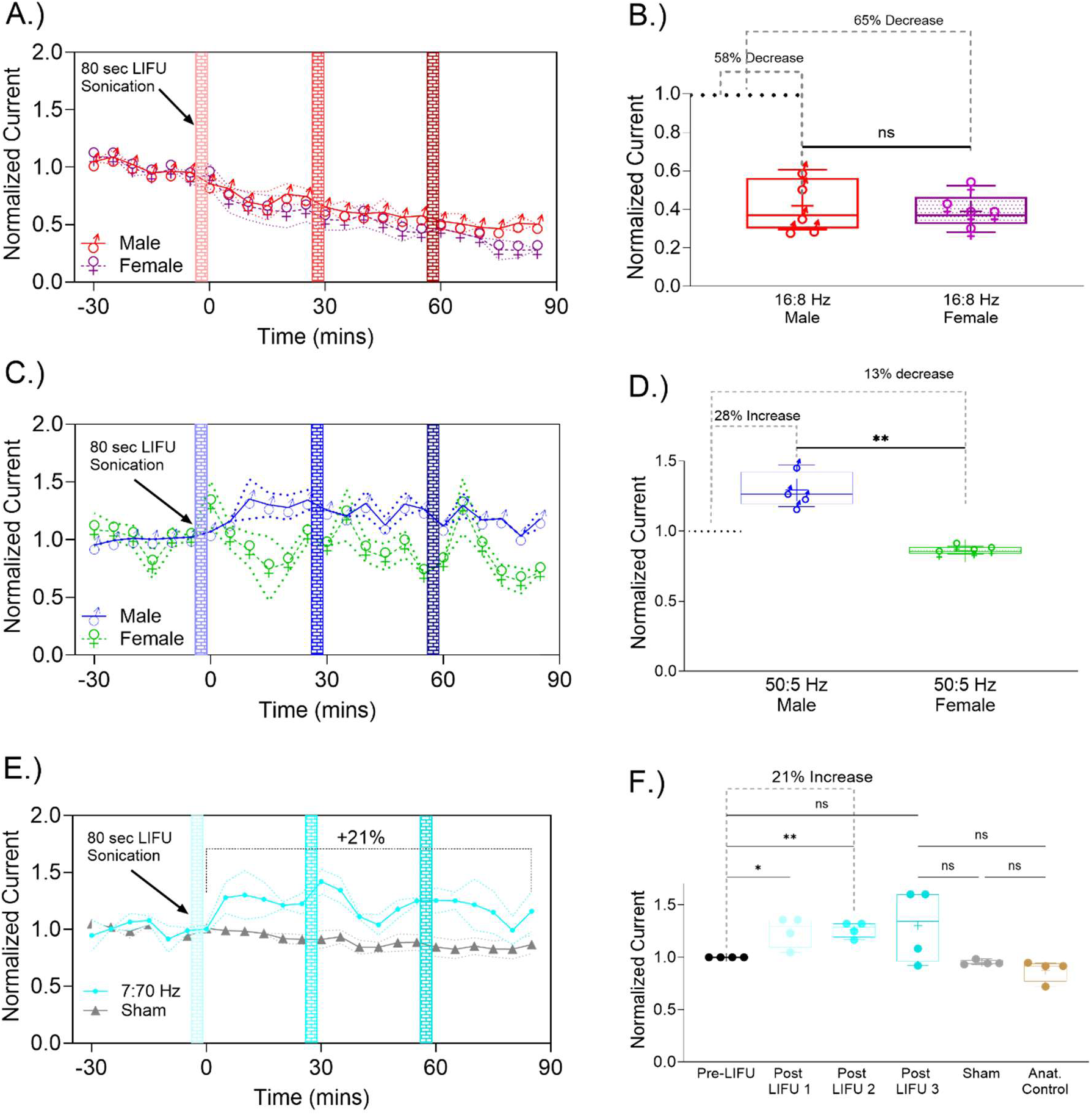
Sex-specific modulation of dopamine release by LIFU. **(A)** Time course of the normalized dopamine current (mean ± SEM) in male (red) and female (purple) rats following 8:16 Hz LIFU. The red bars denote the 80-second stimulation period. Both males and females exhibited a significant decline over time, with males reaching 40.4% of baseline (Phase 4: M = 0.404 ± 0.043) and females reaching 39% (M = 0.390 ± 0.047). Multivariate analysis confirmed significant phase effects for BT (*Pillai’s trace* = 0.929, *F*_(3,14)_ = 60.86, *p* < 0.001) and female BT (*Pillai’s trace* = 0.922, *F*_(3,14)_ = 55.40, *p* < 0.001). **(B)** Boxplots of normalized dopamine levels across experimental conditions. The final-phase (Phase 4) dopamine currents did not differ between the sexes in the 8:16 Hz condition (BT vs. female BT: Δ = 0.015 ± 0.064, *p* = 1.000). Between-group comparisons revealed no significant differences between the males and females at any phase (Phase 2: Δ = 0.002 ± 0.088, *p* = 1.000; Phase 3: Δ = 0.040 ± 0.062, *p* = 0.989; Phase 4: Δ = 0.015 ± 0.064, *p* = 1.000). **(C)** Time course for the 5:50 Hz condition. There was a robust and sustained increase in the number of males (blue) from baseline (Phase 4: M = 1.265 ± 0.047), while the number of females (green) decreased modestly (M = 0.860 ± 0.053). A significant phase effect was detected for *Pillai’s trace* = 0.724, *F*_(3,14)_ = 12.27, *p* < 0.001 but not for HGT (*Pillai’s trace* = 0.392, *F*_(3,14)_ = 3.01, *p* = 0.066). Among the males, dopamine levels significantly increased from baseline to Phase 2 (Δ = –0.302 ± 0.065, *p* = 0.002), Phase 3 (Δ = –0.284 ± 0.046, *p* < 0.001), and Phase 4 (Δ = –0.265 ± 0.047, *p* < 0.001). No within-group differences across phases were significant for HGT (all *p* > 0.10). **(D)** Boxplots of normalized dopamine levels across experimental conditions. In Phase 4, male dopamine levels significantly exceeded female dopamine levels (Δ = 0.405 ± 0.071, *p* < 0.001). **E)** Time course of the normalized dopamine current in females following 7:70 Hz LIFU (cyan) versus sham controls (gray). The 7:70 Hz protocol induces a robust and sustained 21% increase in dopamine release, persisting across all stimulation phases and at 90 minutes poststimulation. **(F)** Boxplot summary of the normalized dopamine current in females across sequential LIFU phases (pre-LIFU, post-LIFU 1–3), anatomical controls, and sham controls. Significant progressive increases were observed following 7:70 Hz LIFU (*p < 0.05, **p < 0.01), but there were no significant changes in the control groups.

In contrast, the 5:50 Hz stimulation paradigm revealed a pronounced difference in dopamine responses between the sexes (Fig. 5C). A sustained increase in dopamine release lasted more than 30 minutes in males, with significant increases observed across all stimulation phases. Multivariate analysis confirmed a significant, phase-dependent increase in dopamine current in males, reflecting consistent and prolonged dopamine enhancement. Only females exhibited a transient increase in dopamine release, which returned to baseline within 10 minutes post-LIFU. Moreover, a significant sex-by-phase difference according to treatment was observed in phases 3 and 4 following the second and third LIFU treatments, respectively, in which females were included. The phase-dependent changes in dopamine current were not statistically significant in females, indicating that their response was brief and did not mirror the sustained enhancement observed in males. Boxplot analysis of Phase 4 dopamine release further confirmed a significant sex-specific difference in this frequency (Fig. 5D), with males showing a 28% increase and females showing a 13% decrease.

To further characterize sex dependent, frequency-specific effects, we examined the impact of theta-high-gamma (7:70 Hz) LIFU stimulation on dopamine release in females (Fig. 5E). This protocol produced a robust and sustained increase in dopamine release (+21%) compared to that of the sham controls, which maintained baseline levels throughout the experiment. The enhanced dopamine response persisted across all three stimulation phases and remained elevated at 90 minutes, indicating that female rats require a greater frequency of increased dopamine release. In males, however, 7:70 Hz LIFU only caused an increase in dopamine after the first stimulation Figure S6). Dopamine levels returned to base line following the second and third LIFU treatments

Figure 5F shows comprehensive statistical comparisons across sequential post-LIFU phases, revealing significant progressive enhancement from baseline (pre-LIFU) through successive stimulations (post-LIFU 1–3). Statistical analysis demonstrated significant differences post-LIFU 1 (p<0.0442) and post-LIFU 2 (p<0.0057) compared to pre-LIFU dopamine release, suggesting a cumulative effect of repeated LIFU stimulation at specific frequencies.

These results demonstrated that while both sexes exhibited comparable inhibition of dopamine release following 8:16 Hz LIFU, only males showed sustained dopamine enhancement at 5:50 Hz. In females, a higher gamma frequency is required to elicit a robust increase in dopamine release. This finding underscores the frequency dependent, sex-specific modulation of dopamine signaling by LIFU and identifies critical parameters for targeted neuromodulation applications.

### Region- and Circuit-Specific Effects of LIFU Stimulation on Neuronal Activity and Astrocytic Response in the PFC and Downstream Signaling in the NAc

Figure 6 illustrates the LIFU-induced modulation of neuronal and astrocytic markers in the PLC and its downstream projection to the NAcc in males. Representative coronal sections demarcating the PLC (Fig. 6A) and NAcc (Fig. 6C) display DAPI, NeuN, cFOS, and GFAP expression. Corresponding immunofluorescence images under sham, 8:16 Hz, and 5:50 Hz conditions are shown in Fig. 6B and D. In both the PLC and NAcc, overall NeuN density and expression remained statistically unchanged across conditions (Fig. 6E, F), and histological assessment revealed no overt cell death (Figure S3).

**Figure 6:**
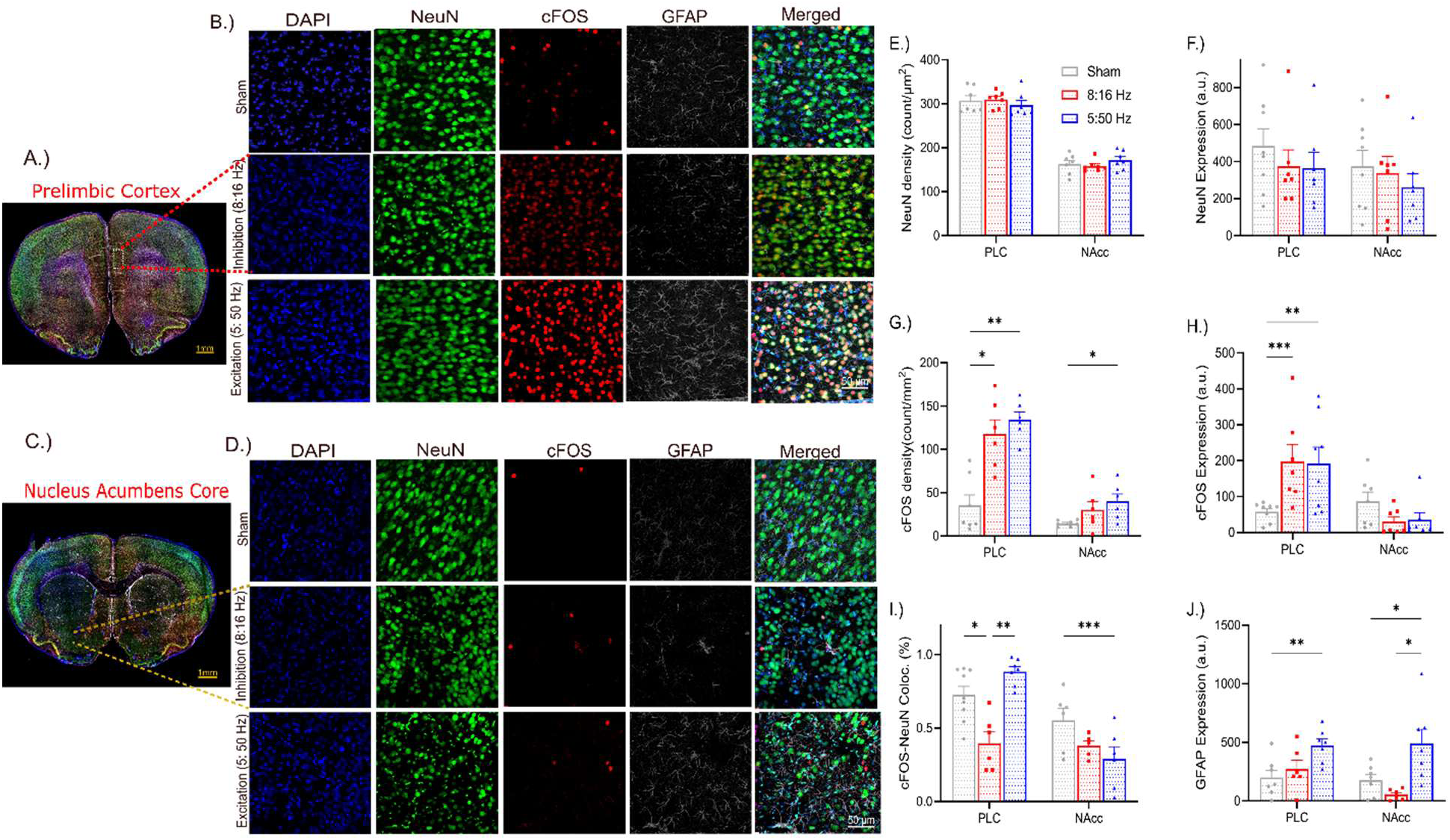
*Effects of* low-intensity focused ultrasound (LIFU) stimulation on neuronal activity and the astrocytic response in the prelimbic cortex (PLC) and downstream signaling in the nucleus accumbens (NAc). A) Anatomical localization of the PLC: Representative coronal brain section illustrating the PLC region targeted for analysis. (B) Immunofluorescence staining for DAPI (nuclear marker, blue), NeuN (neuronal marker, green), cFOS (neuronal activation marker, red), and GFAP (astrocyte marker, white) in the PLC under three experimental conditions: sham control, LIFU at 8:16 Hz, and LIFU at 5:50 Hz. (C) Anatomical localization of the NAcc: Representative coronal brain section depicting the NAcc region targeted for analysis. (D) Immunofluorescence staining of the NAcc under the same three experimental conditions as in (B). (E) Quantitative analysis of NeuN density in the PLC and NAcc across all the experimental conditions. Statistical analysis revealed no significant differences between the groups (p= 0.0954). (F) Quantitative analysis of NeuN expression in the PLC and NAcc across all the experimental conditions. Statistical analysis revealed no significant differences between the groups (p = 0.4338). (G) Quantitative analysis of cFOS density in the PLC and NAcc. In the PLC, both the 8:16 Hz and 5:50 Hz LIFU groups had significantly greater cFOS density than did the sham group, with no significant difference between the two LIFU frequencies (PLC; 8:16 Hz (p=0.01) and 5:50 Hz (p=0.02)). In the NAcc, only the 5:50 Hz group exhibited a significant increase in cFOS density (p=0.03). (H) Quantitative analysis of cFOS expression in the PLC and NAcc. In the PLC, cFOS fluorescence was significantly greater in both LIFU groups than in the sham (5:50 Hz; p<0.0001, 8:16 Hz; p<0.01) group, while no significant differences were observed in the NAcc. (I) Quantitative assessment of colocalization of cFOS+/NeuN+ cells in the PLC and NAcc. In the PLC, the 5:50 Hz group exhibited a significantly greater proportion of cFOS+/NeuN+ cells than the 8:16 Hz group did (p=0.0047). In the NAcc, the 5:50 Hz group had a significantly lower proportion of cFOS+/NeuN+ cells than the sham group (p=0013). (J) Quantitative analysis of GFAP expression in the PLC and NAcc. In both regions, GFAP fluorescence was significantly greater in the 5:50 Hz group than in both the sham and 8:16 Hz groups (PLC: 8:16 Hz; p=0.0159, Sham; p=0.0401; NAcc: 8:16 Hz; p=0.0380, Sham; p=0.0328).

Markers of cellular activation were assessed by cFOS density (number of cFOS+ nuclei per area; Fig. 6G) and total cFOS fluorescence (Fig. 6H). In the PLC, both the 8:16 Hz and 5:50 Hz LIFU groups had significantly greater cFOS density than did the sham group, with no significant difference between the 5:50 Hz and 8:16 Hz groups. Only the 5:50 Hz group exhibited significantly increased cFOS density in the NAcc. cFOS fluorescence significantly increased the PLC in both LIFU groups, while the three groups did not significantly differ from each other in the NAcc.

To further understand the source of the increase in cFOS density observed in both LIFU treatment paradigms, colocalization of cFOS with NeuN was quantified (Fig. 6I). In the PLC, the 5:50 Hz group exhibited a significantly greater proportion of cFOS+/NeuN+ cells than did the 8:16 Hz group, confirming neuronal activation. In the NAcc, however, the 5:50 Hz group showed a significantly lower cFOS+/NeuN+ proportion, indicating that a substantial component of the cFOS signal arose from nonneuronal cells. Consistent with this, GFAP fluorescence—an index of astrocyte activation—was significantly increased after 5:50 Hz treatment compared with after 8:16 Hz treatment in the PLC and after both 8:16 Hz treatment and sham treatment in the NAcc (Fig. 6J).

Finally, to assess circuit specificity, we evaluated cFOS and GFAP in the infralimbic cortex (ILC) and nucleus accumbens shell (NAcs; Supplementary Fig. S4). Although the cFOS density was greater in the LIFU group than in the sham group, GFAP expression remained stable. Moreover, 8:16 Hz LIFU modestly reduced the proportion of cFOS+/NeuN+ cells among both ILCs and NAcs. Collectively, these data demonstrate region- and frequency-specific modulation of neuronal and astrocytic activity by LIFU without affecting cell survival and underscore its potential for targeted neuromodulation of defined frontostriatal circuits.

## Discussion

This study demonstrated that LIFU applied to the PFC can directly influence downstream mesolimbic dopamine release in a frequency-dependent manner. We hypothesized that theta-gamma coupling amplification would promote neuronal synchronization and increase downstream dopamine release in the mesolimbic circuit, whereas theta-beta coupling would have an inhibitory effect. These hypotheses are strongly supported by the observed 28% increase in dopamine release under 5:50 Hz theta-gamma stimulation and the 58% decrease under 16:8 Hz theta-beta stimulation in male rats. Both males and females exhibited comparable dopamine inhibition at 16:8 Hz, but only males exhibited sustained dopamine enhancement at 5:50 Hz, while females were excited at higher gamma frequencies of 70:7 Hz. Moreover, 16:8 Hz and 5:50 Hz LIFU increased neuronal activation in the PLC in males, while 5:50 Hz also upregulated the astrocytic marker GFAP. These results demonstrated that LIFU neuromodulation can mimic neural oscillations and induce inhibitory or excitatory neuromodulation of mesolimbic dopamine signaling via distinct sex- and cell type-specific responses.

The observed frequency-specific effects of LIFU align with established models of neural oscillation function. Beta oscillations mediate inhibitory control by suppressing extraneous neural activity to maintain cognitive stability.^20^ In the PFC, these oscillations regulate working memory through distinct phase-dependent mechanisms.^18,30^ Moreover, the recruitment of the right frontal beta to actively suppress memory retrieval and induce cognitive stopping has been previously reported.^19^ The observed dopamine inhibition with 8:16 Hz and 2:16 Hz LIFU further corroborates this framework. Therefore, our findings quantitatively support previous work, including that of Engel & Fries’ (2010) model, where beta oscillations are proposed to maintain cognitive states through network-level inhibition.^31^

Gamma oscillations, known to facilitate synaptic plasticity and neurotransmitter release through precise spike-timing coordination, are associated with enhanced learning and decision-making.^21,32–34^ Moreover, Zeng *et al*. (2021) showed that theta burst LIFU (5:50 Hz) produced a consistent increase in corticospinal excitability, aligning with the findings of previous studies that highlighted the collaborative role of gamma oscillations with theta rhythms in information encoding.^35,36^ Building upon these observations, we provide further evidence that externally applied oscillatory patterns, delivered via LIFU, can effectively entrain and modulate neurotransmitter systems. Therefore, the observed 28% increase in dopamine density under theta-gamma (5:50 Hz) coupling resonates with established models of cross-frequency coupling mechanisms and supports the role of theta-gamma coupling in facilitating synaptic potentiation.

The use of theta coupling in our study serves dual purposes. First, the theta component helps reduce the overall amount of LIFU delivered, potentially mitigating heat-related concerns while still harnessing the fundamental frequency effects. Second, it leverages the established role of theta interactions in facilitating synaptic potentiation.^16^ While gamma and beta frequency bands have been observed to play specific roles, cross-frequency coupling with hippocampal theta oscillations is important in memory formation and long-term potentiation.^15,16^ Therefore, coupling gamma and beta frequency bands with theta bands is intuitively expected to enhance the long-term potentiation of their noncoupled effect, as observed in this study.

The effects of LIFU were also cell type specific. In the PLC, while the 8:16 Hz LIFU-mediated inhibition of dopamine and 5:50 Hz led to increased dopamine release, both treatments elicited a net increase in neuronal activation, as indicated by heightened cFOS expression and density. This apparent contradiction may reflect preferential activation of inhibitory neurons by 8:16 Hz LIFU, leading to downstream inhibition of dopamine release in the NAcc (Figure 1), suggesting that different cell types within the PLC may respond differently to LIFU stimulation.

In contrast, 5:50 Hz LIFU reveals a fundamentally different profile of cellular engagement. While the colocalization of cFOS with NeuN was markedly greater in the PLC in the 5:50 Hz LIFU group than in the 8:16 Hz group, both groups exhibited lower colocalization in the NAcc than in the Sham group. This observation points to a sizable nonneuronal cFOS source in the 5:50 Hz group. Indeed, only at 5:50 Hz did we observe pronounced upregulation of GFAP in the PLC and NAcc, implicating astrocytes as key drivers of the enhanced cFOS signal. Astrocytes can profoundly influence synaptic plasticity through their high-affinity glutamate transporters and capacity to release gliotransmitters.^37–39^ We therefore propose that 5:50 Hz LIFU recruits astrocytic networks, leading to glutamate-mediated activation of presynaptic NMDA receptors on dopaminergic terminals, ultimately leading to increased NAcc dopamine. ^40–42^

Extending our analysis to ILCs and NAcs revealed a similarly nuanced interplay of frequency- and region-specific effects (Fig. S4). Both the 8:16 and 5:50 Hz paradigms drive a significant increase in cFOS. However, GFAP expression did not change in either region, highlighting the contribution of latent astrocytes. In ILCs, 8:16 Hz suppresses NeuN expression. In the NAcs, the NeuN intensity was significantly dampened only by 8:16 Hz LIFU. While the LIFU-induced modulation of DA release is highly circuit specific (PLC→NAc core), similar levels of activity were observed in ILCs post LIFU. Regional specificity is rather significant in the downstream accumbal regions. We propose that this observation is due to a lower acoustic pressure (40%) reaching the ILC (Figure S1), as shown in our previous study using a similar LIFU intensity. This finding suggested that while functional output (dopamine release) is highly targeted, underlying cellular signaling and stress responses are more broadly spread, likely due to direct or indirect inhibition of neuronal and nonneuronal populations and their projections across neighboring areas.

Previous neuromodulation studies have largely been conducted in male subjects, leaving a critical gap in our understanding of sex-specific neurobiological responses. To address this disparity, we measured dopamine release in male and female rats using three low-intensity focused ultrasound (LIFU) protocols—8:16 Hz, 5:50 Hz, and a high-gamma paradigm—at 7:70 Hz. While 8:16 Hz stimulation produced equivalent dopamine inhibition in both sexes, 5:50 Hz stimulation evoked a robust increase only in males; females showed a modest, nonsignificant decrease indistinguishable from that of sham controls. Mechanistically, the male-specific dopamine boost at 5:50 Hz may reflect stronger synchronization between LIFU-induced gamma–theta entrainment and the intrinsic oscillatory properties of dopaminergic pathways.^43,44^ ^43^ In females, estrogenic modulation of the dopamine system—by increasing dopamine synthesis, release, and D₂ receptor affinity—can introduce nonlinearity or ceiling effects in stimulation-evoked dopamine dynamics.^45–47^ Consequently, such ceiling effects may reduce dopamine release post-LIFU in females. Crucially, only the 7:70 Hz high-gamma protocol succeeded in elevating dopamine release in females, indicating that overcoming ceiling effects to activate female dopaminergic circuits may require a higher gamma frequency parameter.

This divergence aligns with previous reports of distinct sex-specific patterns in theta band processing during spatial navigation observed in EEG studies, suggesting inherent differences in cognitive strategies and neural resource allocation.^48^ Moreover, our findings are supported by several studies demonstrating female sensitivity to high-gamma oscillations. Douton and Carelli (2023) reported pronounced increases in 70–85 Hz LFP power in the nucleus accumbens shell of female rats during conditioned taste aversion.^49^ Theriault *et al*. (2021) observed stress-induced high-gamma enhancements exclusively in females.^50^ More recently, Grablin (2025) showed that 80 Hz optical stimulation of the ILC-NAc pathway modulates innate aversion behaviors only in female rats—reducing aversive responses and increasing neutral behaviors during quinine exposure.^51^ Together, these sex-specific differences in LIFU efficacy suggest fundamental differences in neural resource allocation and cognitive strategy between males and females.

Notably, the temporal dynamics of the LIFU-induced excitatory effects were more robust than those of the inhibitory effects. Excitatory effects (21-28%) were less robust than inhibitory effects on LIFU, lasting only 30 minutes when a one-hour interstimulation interval (ISI) was used under 5:50 Hz LIFU stimulation in males (Figure S5A). This transient and less robust response may reflect an intrinsic neuroprotective mechanism against excitotoxicity. In females, the excitatory effect of 5:50 Hz LIFU was even shorter, lasting only 10 minutes (Figure 5C), and a higher gamma frequency was required to elicit dopamine excitation (Figure 5E). This observation is consistent with sex differences in hormone-dependent baseline striatal activity, dopamine release, and uptake kinetics.^46,52–54^ However, the mechanisms underlying these sex-specific responses remain unclear and warrant further investigation, particularly to assess whether LIFU induces comparable molecular changes in females. A limitation of our study is the lack of estrous cycle monitoring in female rats, given that hormonal fluctuations are known to influence dopamine system function.^55^ Future studies should incorporate estrous cycle tracking to elucidate how hormonal status modulates LIFU-induced neuromodulation.

In conclusion, our findings establish LIFU as a versatile, noninvasive modality capable of frequency- and circuit-specific modulation of dopaminergic signaling, neuronal excitability, and astrocytic engagement within the mesolimbic reward pathway. By systematically varying the pulse repetition frequency, we defined parameter sets that bidirectionally regulate dopamine release and revealed distinct sex-dependent neuromodulatory profiles. This work moves beyond descriptive accounts of baseline sex differences, directly linking sex-specific dopaminergic dynamics to functional outcomes and underscoring the imperative for personalized LIFU protocols. Mechanistically, we elucidated how cross-frequency coupling governs glial–neuronal communication and validated the precision of LIFU in targeting dysregulated dopamine circuits. Translationally, these results lay the groundwork for tailored, sex-informed interventions in psychiatric conditions marked by dopamine imbalance, most notably substance use disorders and major depressive disorder. Future investigations should investigate real-time astrocytic calcium and glutamate fluxes during sonication and evaluate long-term behavioral readouts to refine and optimize the clinical translation of LIFU paradigms.

## Statements and declarations

### Author contributions

Greatness O. Olaitan; Equipment Setup, Experimental verification Investigation and writing; Akhabue K. Okojie; Equipment Setup and Experimental verification; Wendy J. Lynch; Reviewing, B. Jill Venton; Project Administration; Reviewing, Reviewing.

### Conflict of interest

The authors declare no competing interests.

### Ethics Approval

All animal experiments were approved by the University of Virginia Institutional Animal Care and Use Committee (Protocol Number: 3517-10-23).

### Source of biological material

All animals were purchased from Charles River.

### Statement on animal welfare

The animal welfare status was assessed in accordance with the guidelines on animal welfare published by OLAW from the NIH.

### Funding

This project was funded by the National Institutes of Health, National Institute on Drug Abuse, under NIDA 1R01DA052893-01A1

## Notes

### Competing Interest Statement

The authors have declared no competing interest.

