## Supplementary Document for "Frequency-Dependent Modulation of the Prefrontal Cortex by Low-Intensity Focused Ultrasound: Impact on Mesolimbic Dopamine Signaling"

Title: Oscillation frequency-dependent modulation of the prefrontal cortex using Low-Intensity Focused Ultrasound: impact on downstream mesolimbic dopamine.

Authors: Greatness O. Olaitan^a^, Akhabue Kenneth Okojie^b^, Wendy Lynch^b^‬, and B. Jill Venton^a^.‬‬‬‬‬‬‬‬‬‬‬‬‬‬‬‬‬‬‬‬‬‬‬‬‬‬‬‬‬

Affiliations*:*

*Department of Chemistry, University of Virginia, Charlottesville, VA, 22904*

*Psychiatry and Neurobehavioral Sciences, University of Virginia, Charlottesville, VA, USA*

### Ultrasound Characterization

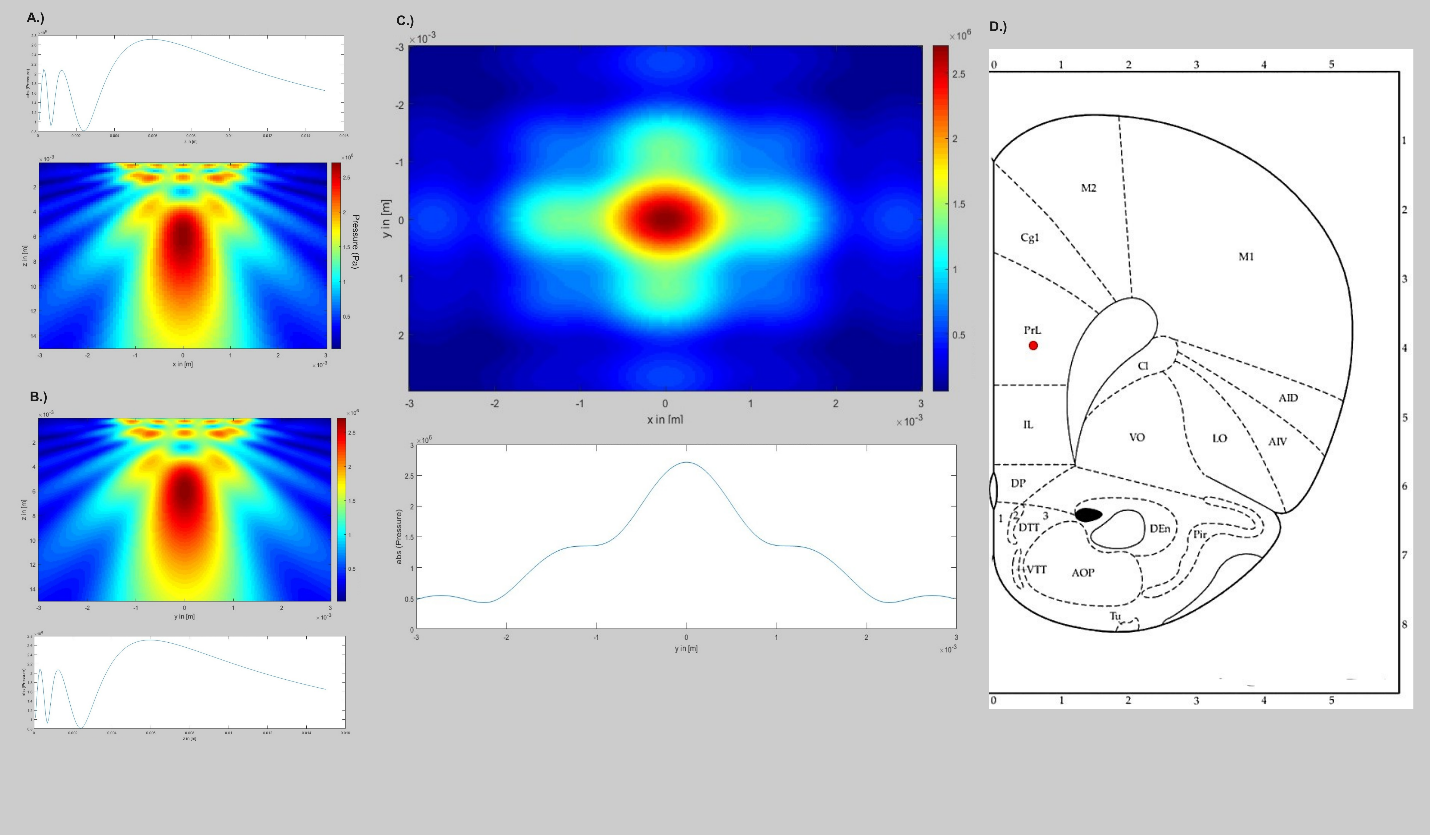

**Figure S1. Acoustic simulation of low-intensity focused ultrasound (LIFU) targeting prefrontal cortical regions.** (**A, B**) Pressure field distributions in the x-z and y-z planes, respectively, demonstrating peak acoustic intensity focused at 5–7 mm from the transducer exit plane. (**C**) Transverse pressure field distribution (x-y plane) showing spatial confinement of the acoustic focus within approximately 0.7 mm diameter. Bottom panel shows the corresponding intensity profile along the focal axis. (**D**) Coronal rat brain section (bregma +3.2 mm) illustrating the anatomical positioning of the prelimbic cortex (PrL) and infralimbic cortex (IL) regions. With the transducer positioned 2 mm above the skull surface, acoustic modeling demonstrates optimal energy delivery to the targeted PrL and IL areas. Color scale indicates normalized acoustic pressure; scale bar represents 1 mm.

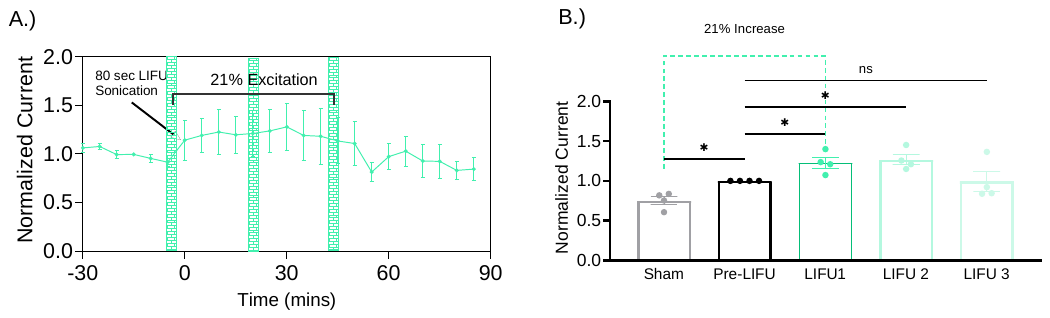

**Figure S2: Gamma Oscillation-dependent excitation of dopamine release by LIFU.** (A) Time course of normalized current following continuous (100% duty cycle) 50 Hz LIFU stimulation. The shaded blue regions indicate 80-second LIFU sonication periods. The graph indicates a 21% increase in normalized current following the first and second stimulation. (B) Quantitative comparison of normalized current across different time points: Bar graph showing normalized current at Sham, Pre-LIFU, LIFU 1, LIFU 2, and LIFU 3 time points. Dopamine release is increased by 21% from Pre-LIFU to Post-LIFU 1 (*), and the increase is persistent following LIFU 2. No significant difference (ns) is observed between LIFU 2 and LIFU 3.

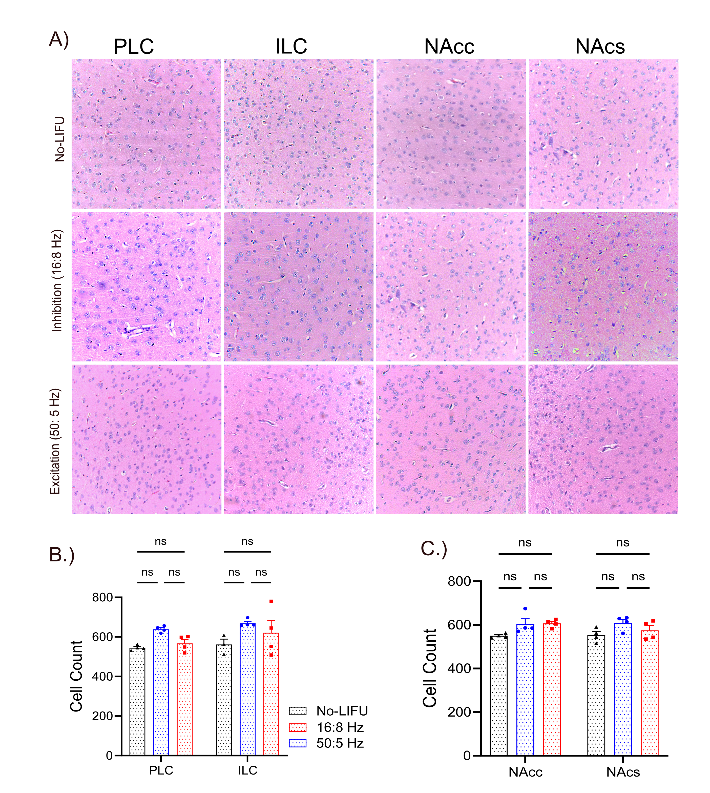

**Figure S3.** ***H&E Staining Shows no Significant Cell Death in LIFU-treated Animals.*** A.) Representative hematoxylin and eosin (H&E)-stained coronal sections of the PLC, ILC, NAcc, and NAcs from animals exposed to no LIFU (top row), LIFU at 16:8 Hz (second row), and LIFU at 50:5 Hz (bottom row). Scale bars = 50 µm. B.) The bar graphs display the quantified expression cell count in the PLC and ILC under each treatment condition compared to the control, showing no significant change. C.) The bar graphs display the quantified expression cell count in the PLC and ILC under each treatment condition compared to the control, showing no significant change. Data are presented as mean ± SEM (n = 4 per group).

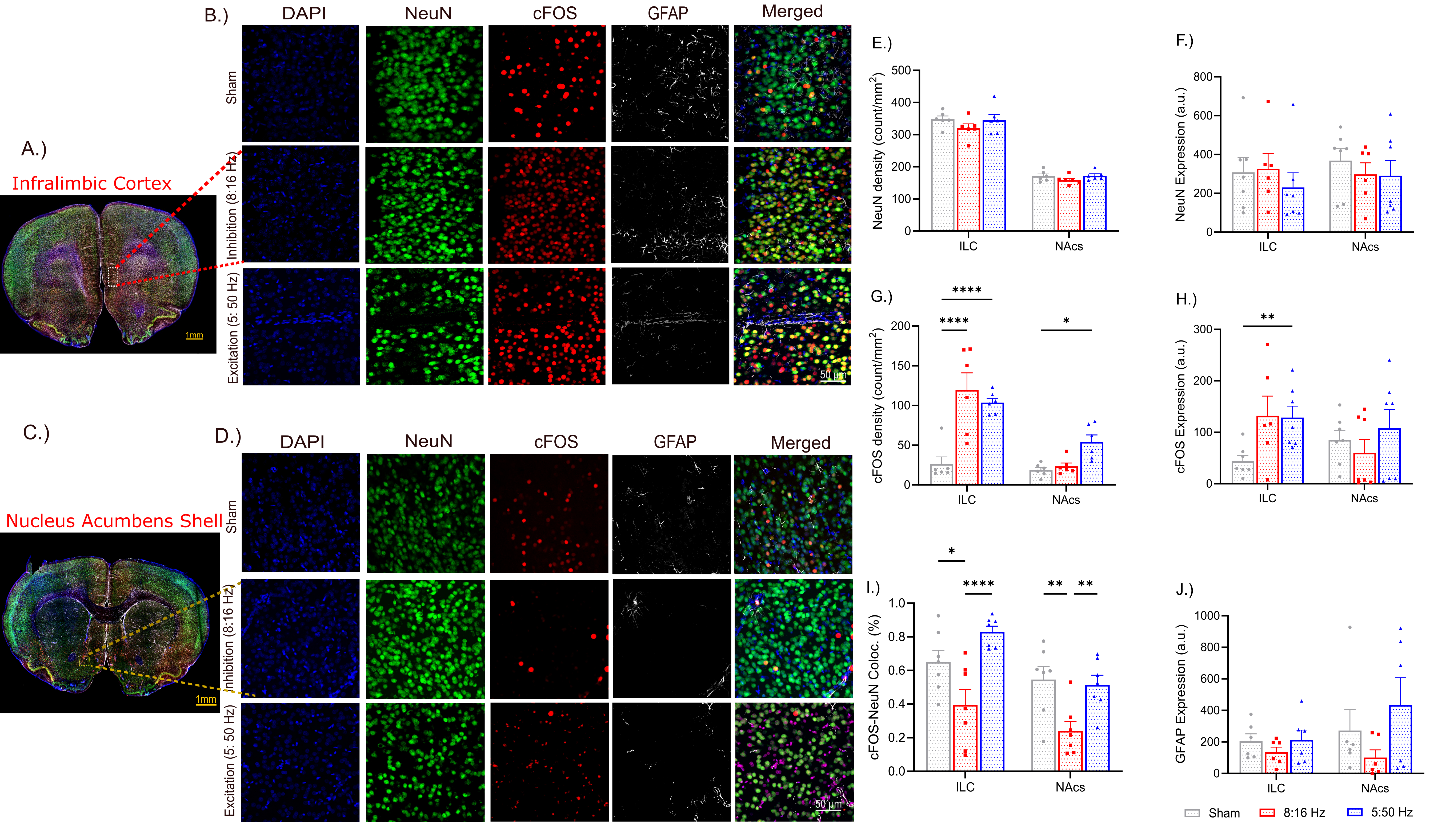

***Figure S4: Effects of Low-Intensity Focused Ultrasound (LIFU) Stimulation on Neuronal Activity and Astrocytic Response in the Infralimbic Cortex (ILC) and Downstream Signaling in the Nucleus Accumbens Shell (NAcs).* (A)** Representative coronal brain section illustrating the ILC and NAcs regions targeted for analysis. (B) Immunofluorescent staining of DAPI (nuclear marker, blue), NeuN (neuronal marker, green), cFOS (neuronal activation marker, red), and GFAP (astrocyte marker, white) in the ILC under three experimental conditions: no-LIFU control, LIFU at 16:8 Hz (Inhibition), and LIFU at 50:5 Hz (Excitation). (C) Representative coronal brain section depicting the ILC and NAcs regions targeted for analysis. (D) Immunofluorescent staining in the NAcs under the identical three experimental conditions as in (B). (E) Quantitative analysis of NeuN density in the ILC and NAcs across all experimental conditions. Statistical analysis revealed no significant differences between groups for either region. (F) Quantitative analysis of NeuN expression in the ILC and NAcs across all experimental conditions. Statistical analysis demonstrated no significant differences across all conditions. (G) Quantitative analysis of cFOS density in the ILC and NAcs. LIFU stimulation at 50:5 Hz elicited a significant increase in cFOS density within the ILC compared to no-LIFU controls (p < 0.0001) and LIFU at 16:8 Hz (p < 0.0001). In the NAcs, only the 5:50 Hz group showed a significant increase compared to no-LIFU controls (p = 0.0148). (H) Quantitative analysis of cFOS expression in the ILC and NAcs. The results showed significant increases in the ILC for both LIFU frequencies (16:8 Hz: p = 0.0265, 50:5 Hz: p = 0.0001, F _(2, 20)_ = 5.772) and no significant difference in the NAcs (p = 0.9945). (I) Quantitative assessment of cFOS+/NeuN+ co-localization in the ILC and NAcs expressed as a percentage. LIFU stimulation at 8:16 Hz resulted in a significant decrease in cFOS+/NeuN+ co-localization within the ILC compared to the sham and 5:50 Hz group (Sham: p = 0.0103, 50:5 Hz: p <0.0001, F _(1, 36)_ = 12.36). A similar trend was observed in the NAcs (Sham: p = 0.0027, 50:5 Hz: p = 0.0064, F _(1, 36)_ = 12.36). (J) Quantitative analysis of GFAP expression in the ILC and NAcs. Statistical analysis indicated no significant difference in GFAP expression in the ILC and the NAcs.

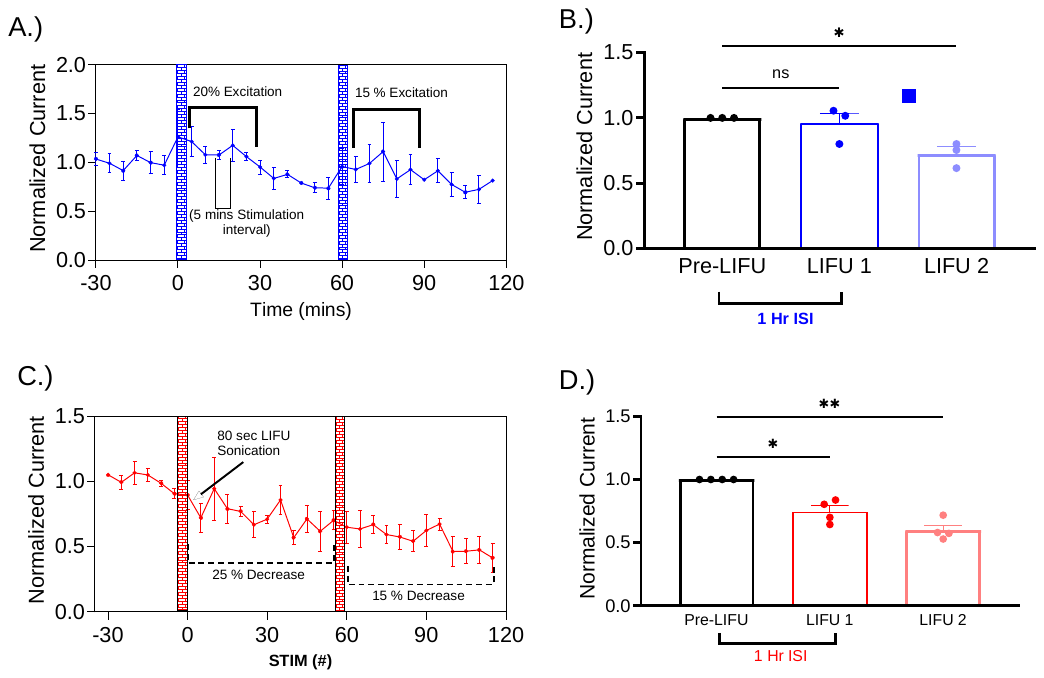

***Figure S5. Temporal dynamics and robustness of LIFU Stimulation with 1-hour Inter-Stimulation Interval (ISI).*** (A) Time course of normalized current following excitatory LIFU stimulation. Blue shaded regions indicate LIFU stimulations. Annotations show 20% and 15% dopamine excitation lasting for only 30 minutes after the respective LIFU stimulation points. (B) Quantitative comparison of normalized current following excitatory LIFU stimulation: Bar graph showing average normalized dopamine oxidation peak current at pre-LIFU, LIFU 1, and LIFU 2 points shows a significant difference (*) between Pre-LIFU and LIFU 2 is indicated. (C) Time course of normalized current following inhibitory LIFU stimulation: Line graph showing normalized current over time (minutes). Red shaded regions indicate LIFU stimulations. Annotations show 25% and 15% dopamine inhibition lasting for 1 hour after the respective LIFU stimulation points. (D) Quantitative comparison of normalized current following inhibitory LIFU stimulation: Bar graph showing average normalized dopamine oxidation peak at pre-LIFU, LIFU 1, and LIFU 2 time points. Significant differences (**) between pre-LIFU and LIFU 1, and LIFU 1 and LIFU 2 are indicated.

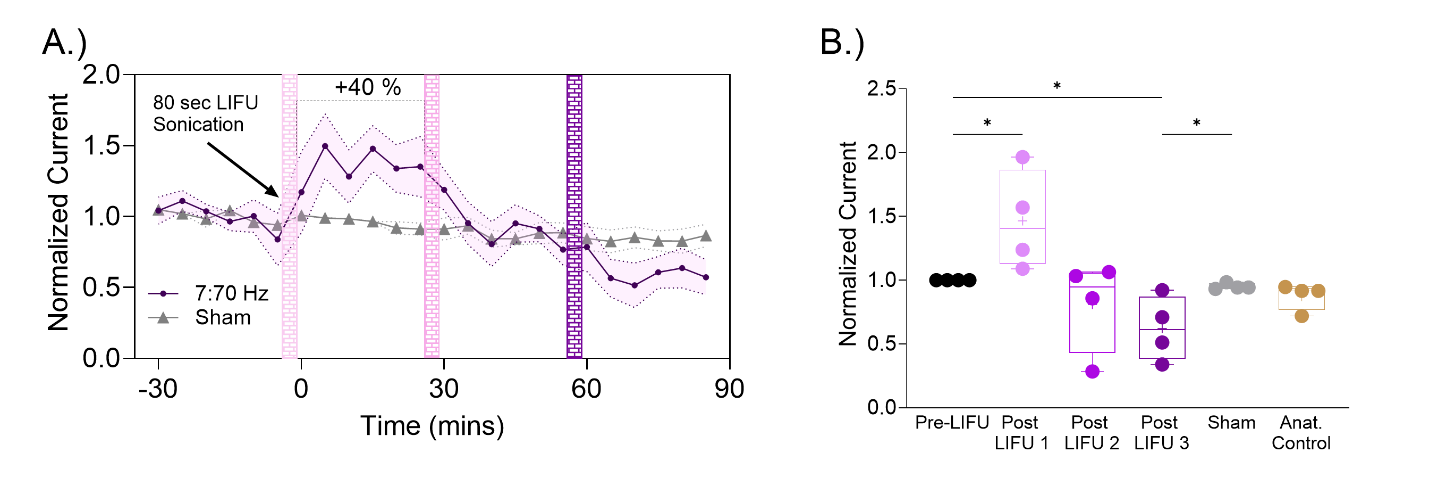

**Figure S6: High Gamma (7:70 Hz) Oscillation-dependent modulation of dopamine release by LIFU in Male rats.** (A) Time course of normalized current following 7:70 Hz LIFU stimulation. The shaded purple regions indicate 80-second LIFU sonication periods. The graph indicates a 40% increase in normalized current following the first stimulation which then returns to baseline upon second LIFU stimulation. (B) Quantitative comparison of normalized current across different time points: Bar graph showing normalized current at Sham, Pre-LIFU, LIFU 1, LIFU 2, and LIFU 3 time points. Dopamine release is increased by 40% from Pre-LIFU to Post-LIFU 1 (*), but there is no significant increase following LIFU 2 or 3. No significant difference (ns) is observed between LIFU 2 and LIFU 3.

**Table S1A:** Between-group comparisons of Normalized DA from Figure 1c.

| **Pairwise Comparisons** | | | | |
| --- | --- | --- | --- | --- |
| (I) Group | (J) Group | Mean Difference (I-J) | Std. Error | Sig.^b^ |
| 16-8 | Anat | -.289^*^ | .066 | .003 |
|  | Sham | -.246^*^ | .066 | .009 |
| Anat | 16-8 | .289^*^ | .066 | .003 |
|  | Sham | .044 | .066 | .890 |
| Sham | 16-8 | .246^*^ | .066 | .009 |
|  | Anat | -.044 | .066 | .890 |

**Table S1B:** Normalized DA across groups and phases in Figure 1c.

| **Estimates** | | | |
| --- | --- | --- | --- |
| Measure: Normalized DA | | | |
| Group | Phase | Mean | Std. Error |
| 16-8 | 1 | 1.000 | .000 |
|  | 2 | .736 | .049 |
|  | 3 | .552 | .084 |
|  | 4 | .419 | .076 |
| Anat | 1 | 1.000 | .000 |
|  | 2 | .942 | .049 |
|  | 3 | .976 | .084 |
|  | 4 | .947 | .076 |
| Sham | 1 | 1.000 | .000 |
|  | 2 | .964 | .049 |
|  | 3 | .884 | .084 |
|  | 4 | .842 | .076 |

**Table S1C:** Post hoc between-group comparisons of Normalized DA at each phase in Figure 1c.

| Measure: Pairwise Comparisons | | | | |  |  |
| --- | --- | --- | --- | --- | --- | --- |
| Measure: Normalized DA | | |  | |  |  |
| Phase | (I) Group | (J) Group | | Mean Difference (I-J) | Std. Error | Sig.b |
| 1 | 16-8 | Anat | 0 | | 0 | . |
|  | 16-8 | Sham | 0 | | 0 | . |
|  | Anat | Sham | 0 | | 0 | . |
| 2 | 16-8 | Anat | -.206* | | 0.069 | 0.033 |
|  | 16-8 | Sham | -.228* | | 0.069 | 0.018 |
|  | Anat | Sham | -0.022 | | 0.069 | 0.985 |
| 3 | 16-8 | Anat | -.424* | | 0.119 | 0.012 |
|  | 16-8 | Sham | -.333* | | 0.119 | 0.048 |
|  | Anat | Sham | 0.091 | | 0.119 | 0.841 |
| 4 | 16-8 | Anat | -.528* | | 0.108 | 0.001 |
|  | 16-8 | Sham | -.423* | | 0.108 | 0.006 |
|  | Anat | Sham | 0.106 | | 0.108 | 0.722 |

**Table S1D:** Post hoc within-group comparisons of Normalized DA across phases from Figure 1c.

| **Group** | **(I) Phase** | **(J) Phase** | **Mean Difference (I-J)** | **Std. Error** | **Sig.b** |
| --- | --- | --- | --- | --- | --- |
| 16-8 | 1 | 2 | .264* | .049 | <.001 |
| 16-8 | 1 | 3 | .448* | .084 | .001 |
| 16-8 | 1 | 4 | .581* | .076 | <.001 |
| 16-8 | 2 | 3 | .185 | .075 | .167 |
| 16-8 | 2 | 4 | .317* | .066 | .003 |
| 16-8 | 3 | 4 | .133* | .029 | .004 |
| Anat | 1 | 2 | .058 | .049 | .828 |
| Anat | 1 | 3 | .024 | .084 | 1.000 |
| Anat | 1 | 4 | .053 | .076 | .985 |
| Anat | 2 | 3 | -.034 | .075 | .998 |
| Anat | 2 | 4 | -.005 | .066 | 1.000 |
| Anat | 3 | 4 | .028 | .029 | .923 |
| Sham | 1 | 2 | .036 | .049 | .979 |
| Sham | 1 | 3 | .116 | .084 | .726 |
| Sham | 1 | 4 | .158 | .076 | .310 |
| Sham | 2 | 3 | .080 | .075 | .891 |
| Sham | 2 | 4 | .123 | .066 | .426 |
| Sham | 3 | 4 | .043 | .029 | .663 |

**Table S1E: Multivariate** tests for the simple effect of Phase within each group in Figures 1b and c.

| **Multivariate Tests** | | | | | | |
| --- | --- | --- | --- | --- | --- | --- |
| Group | | Value | F | Hypothesis df | Error df | Sig. |
| 16-8 | Pillai's trace | .880 | 24.475^a^ | 3.000 | 10.000 | <.001 |
|  | Wilks' lam16:2a | .120 | 24.475^a^ | 3.000 | 10.000 | <.001 |
|  | Hotelling's trace | 7.342 | 24.475^a^ | 3.000 | 10.000 | <.001 |
|  | Roy's largest root | 7.342 | 24.475^a^ | 3.000 | 10.000 | <.001 |
| Anat | Pillai's trace | .167 | .669^a^ | 3.000 | 10.000 | .590 |
|  | Wilks' lam16:2a | .833 | .669^a^ | 3.000 | 10.000 | .590 |
|  | Hotelling's trace | .201 | .669^a^ | 3.000 | 10.000 | .590 |
|  | Roy's largest root | .201 | .669^a^ | 3.000 | 10.000 | .590 |
| Sham | Pillai's trace | .380 | 2.047^a^ | 3.000 | 10.000 | .171 |
|  | Wilks' lam16:2a | .620 | 2.047^a^ | 3.000 | 10.000 | .171 |
|  | Hotelling's trace | .614 | 2.047^a^ | 3.000 | 10.000 | .171 |
|  | Roy's largest root | .614 | 2.047^a^ | 3.000 | 10.000 | .171 |

**Table S2A:** Between-group comparisons of Normalized DA from Figure 2 b.

| (I) Group | (J) Group | Mean Difference (I-J) | Std. Error | Sig.^b^ |
| --- | --- | --- | --- | --- |
| Anat | 16:2 | .218^*^ | .053 | .004 |
| Anat | 16:8 | .285^*^ | .053 | <.001 |
| Anat | Sham | .044 | .055 | .968 |
| 16:2 | 16:8 | .068 | .050 | .727 |
| Sham | 16:2 | .174^*^ | .053 | .023 |
| Sham | 16:8 | .242^*^ | .053 | .001 |

**Table S2B:** Normalized DA across groups and phases in Figure 2 b.

| Group | Phase | Mean | Std. Error |
| --- | --- | --- | --- |
| Anat | 1 | 1.000 | .000 |
|  | 2 | .942 | .046 |
|  | 3 | .976 | .070 |
|  | 4 | .947 | .066 |
| 16:2 | 1 | 1.000 | .000 |
|  | 2 | .826 | .042 |
|  | 3 | .625 | .064 |
|  | 4 | .542 | .060 |
| 16:8 | 1 | 1.000 | .000 |
|  | 2 | .750 | .042 |
|  | 3 | .568 | .064 |
|  | 4 | .404 | .060 |
| Sham | 1 | 1.000 | .000 |
|  | 2 | .964 | .046 |
|  | 3 | .884 | .070 |
|  | 4 | .842 | .066 |

**Table S2C:** Post hoc between-group comparisons of Normalized DA at each phase in Figure 2 b.

| Phase | (I) Group | (J) Group | Mean Difference (I-J) | | Std. Error | | Sig.^b^ |
| --- | --- | --- | --- | --- | --- | --- | --- |
| 1 | Anat | 16:2 | .000 | .000 | | . | |
|  |  | 16:8 | .000 | .000 | | . | |
|  |  | Sham | .000 | .000 | | . | |
|  | 16:2 | Anat | .000 | .000 | | . | |
|  |  | 16:8 | .000 | .000 | | . | |
|  |  | Sham | .000 | .000 | | . | |
|  | 16:8 | Anat | .000 | .000 | | . | |
|  |  | 16:2 | .000 | .000 | | . | |
|  |  | Sham | .000 | .000 | | . | |
|  | Sham | Anat | .000 | .000 | | . | |
|  |  | 16:2 | .000 | .000 | | . | |
|  |  | 16:8 | .000 | .000 | | . | |
| 2 | Anat | 16:2 | .116 | .063 | | .394 | |
|  |  | 16:8 | .191^*^ | .063 | | .040 | |
|  | 16:2 | 16:8 | .075 | .060 | | .780 | |
|  | Sham | Anat | .022 | .065 | | 1.000 | |
|  |  | 16:2 | .138 | .063 | | .218 | |
|  |  | 16:8 | .214^*^ | .063 | | .018 | |
| 3 | Anat | 16:2 | .350^*^ | .095 | | .010 | |
|  |  | 16:8 | .407^*^ | .095 | | .003 | |
|  |  | Sham | .091 | .099 | | .937 | |
|  | 16:2 | 16:8 | .057 | .090 | | .990 | |
|  | Sham | 16:2 | .259 | .095 | | .079 | |
|  |  | 16:8 | .316^*^ | .095 | | .022 | |
| 4 | Anat | 16:2 | .405^*^ | .089 | | .001 | |
|  |  | 16:8 | .543^*^ | .089 | | <.001 | |
|  |  | Sham | .106 | .093 | | .850 | |
|  | 16:2 | 16:8 | .138 | .085 | | .544 | |
|  | Sham | 16:2 | .300^*^ | .089 | | .021 | |
|  |  | 16:8 | .437^*^ | .089 | | <.001 | |

**Table S2D:** Post hoc within-group comparisons of Normalized DA across phases from Figure 2 b.

| Group | (I) Phase | (J) Phase | Mean Difference (I-J) | Std. Error | Sig.^b^ |
| --- | --- | --- | --- | --- | --- |
| Anat | 1 | 2 | 0.058 | 0.046 | 0.782 |
|  | 1 | 3 | 0.024 | 0.07 | 1 |
|  | 1 | 4 | 0.053 | 0.066 | 0.967 |
|  | 2 | 3 | -0.034 | 0.065 | 0.996 |
|  | 2 | 4 | -0.005 | 0.067 | 1 |
|  | 3 | 4 | 0.028 | 0.035 | 0.966 |
| 16:2 | 1 | 2 | .174^*^ | 0.042 | 0.004 |
|  | 1 | 3 | .375^*^ | 0.064 | <.001 |
|  | 1 | 4 | .458^*^ | 0.06 | <.001 |
|  | 2 | 1 | -.174^*^ | 0.042 | 0.004 |
|  | 2 | 3 | .200^*^ | 0.059 | 0.02 |
|  | 2 | 4 | .284^*^ | 0.061 | 0.001 |
|  | 3 | 1 | -.375^*^ | 0.064 | <.001 |
|  | 3 | 2 | -.200^*^ | 0.059 | 0.02 |
|  | 3 | 4 | 0.083 | 0.032 | 0.103 |
|  | 4 | 1 | -.458^*^ | 0.06 | <.001 |
|  | 4 | 2 | -.284^*^ | 0.061 | 0.001 |
|  | 4 | 3 | -0.083 | 0.032 | 0.103 |
| 16:8 | 1 | 2 | .250^*^ | 0.042 | <.001 |
|  | 1 | 3 | .432^*^ | 0.064 | <.001 |
|  | 1 | 4 | .596^*^ | 0.06 | <.001 |
|  | 2 | 1 | -.250^*^ | 0.042 | <.001 |
|  | 2 | 3 | .182^*^ | 0.059 | 0.038 |
|  | 2 | 4 | .346^*^ | 0.061 | <.001 |
|  | 3 | 1 | -.432^*^ | 0.064 | <.001 |
|  | 3 | 2 | -.182^*^ | 0.059 | 0.038 |

**Table S2E:: Multivariate** tests for the simple effect of Phase within each group in Figures 2 b and c.

| **Multivariate Tests** | | | | | | |
| --- | --- | --- | --- | --- | --- | --- |
| Group | | Value | F | Hypothesis df | Error df | Sig. |
| Anat | Pillai's trace | .141 | .873^a^ | 3.000 | 16.000 | .475 |
|  | Wilks' lambda | .859 | .873^a^ | 3.000 | 16.000 | .475 |
|  | Hotelling's trace | .164 | .873^a^ | 3.000 | 16.000 | .475 |
| 16:2 | Pillai's trace | .785 | 19.512^a^ | 3.000 | 16.000 | <.001 |
|  | Wilks' lambda | .215 | 19.512^a^ | 3.000 | 16.000 | <.001 |
|  | Hotelling's trace | 3.659 | 19.512^a^ | 3.000 | 16.000 | <.001 |
| 16:8 | Pillai's trace | .878 | 38.473^a^ | 3.000 | 16.000 | <.001 |
|  | Wilks' lambda | .122 | 38.473^a^ | 3.000 | 16.000 | <.001 |
|  | Hotelling's trace | 7.214 | 38.473^a^ | 3.000 | 16.000 | <.001 |
| Sham | Pillai's trace | .270 | 1.968^a^ | 3.000 | 16.000 | .160 |
|  | Wilks' lambda | .730 | 1.968^a^ | 3.000 | 16.000 | .160 |
|  | Hotelling's trace | .369 | 1.968^a^ | 3.000 | 16.000 | .160 |

**Table S3A:** Between-group comparisons of Normalized DA from Figure 3 b.

**Pairwise Comparisons**

| **(I) Group** | **(J) Group** | **Mean Difference (I-J)** | **Std. Error** | **Sig.b** |
| --- | --- | --- | --- | --- |
| 50:5 | Anat | .247* | .063 | .006 |
| 50:5 | Sham | .290* | .063 | .002 |
| Anat | Sham | .044 | .063 | .874 |

**Table S3B:** Normalized DA across groups and phases in Figure 3 b.

| **Estimates** | | | |
| --- | --- | --- | --- |
| Measure: Normalized DA | | | |
| Group | Phase | Mean | Std. Error |
| Anat | 1 | 1.000 | .000 |
|  | 2 | .942 | .052 |
|  | 3 | .976 | .080 |
|  | 4 | .947 | .075 |
| 50:5 | 1 | 1.000 | .000 |
|  | 2 | 1.302 | .052 |
|  | 3 | 1.284 | .080 |
|  | 4 | 1.265 | .075 |
| Sham | 1 | 1.000 | .000 |
|  | 2 | .964 | .052 |
|  | 3 | .884 | .080 |
|  | 4 | .842 | .075 |

**Table S3C:** Post hoc between-group comparisons of Normalized DA at each phase in Figure 3 b.

**Pairwise Comparisons**

Measure: Normalized DA

| **Phase** | **(I) Group** | **(J) Group** | **Mean Difference (I-J)** | **Std. Error** | **Sig.b** |
| --- | --- | --- | --- | --- | --- |
| 1 | Anat | 50:5 | .000 | .000 | . |
| 1 | Anat | Sham | .000 | .000 | . |
| 1 | 50:5 | Sham | .000 | .000 | . |
| 2 | 50:5 | Anat | .360* | .074 | .001 |
| 2 | 50:5 | Sham | .338* | .074 | .002 |
| 2 | Sham | Anat | .022 | .074 | .988 |
| 3 | 50:5 | Anat | .308 | .113 | .055 |
| 3 | 50:5 | Sham | .400* | .113 | .013 |
| 3 | Anat | Sham | .091 | .113 | .822 |
| 4 | 50:5 | Anat | .318* | .106 | .033 |
| 4 | 50:5 | Sham | .424* | .106 | .005 |
| 4 | Anat | Sham | .106 | .106 | .711 |

**Table S3D:** Post hoc within-group comparisons of Normalized DA across phases from Figure 3 b and c.

**Pairwise Comparisons**

Measure: Normalized DA

| **Group** | **Phase 1** | **Phase 2** | **Mean Diff (I-J)** | **Std. Error** | **Sig.b** |
| --- | --- | --- | --- | --- | --- |
| Anat | 1 | 2 | 0.058 | 0.052 | .871 |
| Anat | 1 | 3 | 0.024 | 0.080 | 1.000 |
| Anat | 1 | 4 | 0.053 | 0.075 | .983 |
| Anat | 3 | 2 | 0.034 | 0.080 | .999 |
| Anat | 4 | 2 | 0.005 | 0.076 | 1.000 |
| Anat | 3 | 4 | 0.028 | 0.033 | .954 |
| 50:5 | 2 | 1 | 0.302* | 0.052 | <.001 |
| 50:5 | 3 | 1 | 0.284* | 0.080 | .024 |
| 50:5 | 4 | 1 | 0.265* | 0.075 | .024 |
| 50:5 | 2 | 3 | 0.018 | 0.080 | 1.000 |
| 50:5 | 2 | 4 | 0.037 | 0.076 | .998 |
| 50:5 | 4 | 3 | 0.019 | 0.033 | .994 |
| Sham | 1 | 2 | 0.036 | 0.052 | .986 |
| Sham | 1 | 3 | 0.116 | 0.080 | .685 |
| Sham | 1 | 4 | 0.158 | 0.075 | .293 |
| Sham | 2 | 3 | 0.080 | 0.080 | .914 |
| Sham | 2 | 4 | 0.123 | 0.076 | .570 |
| Sham | 3 | 4 | 0.043 | 0.033 | .765 |

**Table S3E: Multivariate** tests for the simple effect of Phase within each group in Figures 1b and c.

| **Multivariate Tests** | | | | | |  | | |
| --- | --- | --- | --- | --- | --- | --- | --- | --- |
| Group | | Value | F | Hypothesis df | | Error df | | Sig. |
| Anat | Pillai's trace | .154 | .608^a^ | 3.000 | 10.000 | | .625 | |
|  | Wilks' lam16:2a | .846 | .608^a^ | 3.000 | 10.000 | | .625 | |
|  | Hotelling's trace | .182 | .608^a^ | 3.000 | 10.000 | | .625 | |
|  | Roy's largest root | .182 | .608^a^ | 3.000 | 10.000 | | .625 | |
| 50:5 | Pillai's trace | .751 | 10.060^a^ | 3.000 | 10.000 | | .002 | |
|  | Wilks' lam16:2a | .249 | 10.060^a^ | 3.000 | 10.000 | | .002 | |
|  | Hotelling's trace | 3.018 | 10.060^a^ | 3.000 | 10.000 | | .002 | |
|  | Roy's largest root | 3.018 | 10.060^a^ | 3.000 | 10.000 | | .002 | |
| Sham | Pillai's trace | .331 | 1.648^a^ | 3.000 | 10.000 | | .240 | |
|  | Wilks' lam16:2a | .669 | 1.648^a^ | 3.000 | 10.000 | | .240 | |
|  | Hotelling's trace | .494 | 1.648^a^ | 3.000 | 10.000 | | .240 | |
|  | Roy's largest root | .494 | 1.648^a^ | 3.000 | 10.000 | | .240 | |

**Table S4A:** Between-group comparisons of Normalized DA from Figure 4 b.

**Pairwise Comparisons**

Measure: Normalized DA

| **Group I** | **Group J** | **Mean Diff (I-J)** | **Std. Error** | **Sig.b** |
| --- | --- | --- | --- | --- |
| 50:5 | Anat | 0.247* | 0.081 | .047 |
| 50:5 | 28:7 | 0.266* | 0.081 | .029 |
| 50:5 | Sham | 0.290* | 0.081 | .015 |
| Anat | 28:7 | 0.019 | 0.081 | 1.000 |
| Anat | Sham | 0.044 | 0.081 | .996 |
| 28:7 | Sham | 0.024 | 0.081 | 1.000 |

**Table S4B:** Normalized DA across groups and phases in Figure 4 b.

| **Estimates** | | | | | |
| --- | --- | --- | --- | --- | --- |
| Measure: Normalized DA | | | | | |
| Group | Phase | | Mean | | Std. Error |
| Anat | 1 | 1.000 | | .000 | |
|  | 2 | .942 | | .070 | |
|  | 3 | .976 | | .091 | |
|  | 4 | .947 | | .089 | |
| 50:5 | 1 | 1.000 | | .000 | |
|  | 2 | 1.302 | | .070 | |
|  | 3 | 1.284 | | .091 | |
|  | 4 | 1.265 | | .089 | |
| 28:7 | 1 | 1.000 | | .000 | |
|  | 2 | .982 | | .070 | |
|  | 3 | .944 | | .091 | |
|  | 4 | .861 | | .089 | |
| Sham | 1 | 1.000 | | .000 | |
|  | 2 | .964 | | .070 | |
|  | 3 | .884 | | .091 | |
|  | 4 | .842 | | .089 | |

**Table S4C:** Post hoc between-group comparisons of Normalized DA at each phase in Figure 4 b.

**Pairwise Comparisons**

Measure: Normalized DA

| **Phase** | **Group I** | **Group J** | **Mean Diff (I-J)** | **Std. Error** | **Sig.b** |
| --- | --- | --- | --- | --- | --- |
| 1 | Anat | 50:5 | 0.000 | 0.000 | . |
| 1 | Anat | 28:7 | 0.000 | 0.000 | . |
| 1 | Anat | Sham | 0.000 | 0.000 | . |
| 1 | 50:5 | 28:7 | 0.000 | 0.000 | . |
| 1 | 50:5 | Sham | 0.000 | 0.000 | . |
| 1 | 28:7 | Sham | 0.000 | 0.000 | . |
| 2 | 50:5 | Anat | 0.360* | 0.099 | .013 |
| 2 | 50:5 | 28:7 | 0.321* | 0.099 | .031 |
| 2 | 50:5 | Sham | 0.338* | 0.099 | .021 |
| 2 | 28:7 | Anat | 0.040 | 0.099 | .999 |
| 2 | Sham | Anat | 0.022 | 0.099 | 1.000 |
| 2 | Sham | 28:7 | 0.018 | 0.099 | 1.000 |
| 3 | 50:5 | Sham | 0.400* | 0.129 | .041 |
| 3 | 50:5 | Anat | 0.308 | 0.129 | .165 |
| 3 | 50:5 | 28:7 | 0.340 | 0.129 | .104 |
| 3 | Sham | Anat | 0.091 | 0.129 | .982 |
| 3 | 28:7 | Sham | 0.060 | 0.129 | .998 |
| 4 | 50:5 | 28:7 | 0.404* | 0.126 | .033 |
| 4 | 50:5 | Sham | 0.424* | 0.126 | .024 |
| 4 | 50:5 | Anat | 0.318 | 0.126 | .129 |
| 4 | Sham | Anat | 0.106 | 0.126 | .960 |
| 4 | 28:7 | Sham | 0.020 | 0.126 | 1.000 |

**Table S4D:** Post hoc within-group comparisons of Normalized DA across phases from Figure 4 b

**Pairwise Comparisons**

| **Group** | **Phase I** | **Phase J** | **Mean Diff (I-J)** | **Std. Error** | **Sig.b** |
| --- | --- | --- | --- | --- | --- |
| **Anat** | 1 | 2 | 0.058 | 0.070 | .961 |
| Anat | 1 | 3 | 0.024 | 0.091 | 1.000 |
| Anat | 1 | 4 | 0.053 | 0.089 | .993 |
| Anat | 3 | 2 | 0.034 | 0.070 | .998 |
| Anat | 4 | 2 | 0.005 | 0.069 | 1.000 |
| Anat | 3 | 4 | 0.028 | 0.032 | .945 |
| **50:5** | 2 | 1 | 0.302* | 0.070 | .003 |
| 50:5 | 3 | 1 | 0.284* | 0.091 | .040 |
| 50:5 | 4 | 1 | 0.265 | 0.089 | .053 |
| 50:5 | 2 | 3 | 0.018 | 0.070 | 1.000 |
| 50:5 | 2 | 4 | 0.037 | 0.069 | .996 |
| 50:5 | 3 | 4 | 0.019 | 0.032 | .993 |
| **28:7** | 1 | 2 | 0.018 | 0.070 | 1.000 |
| 28:7 | 1 | 3 | 0.056 | 0.091 | .992 |
| 28:7 | 1 | 4 | 0.139 | 0.089 | .594 |
| 28:7 | 2 | 3 | 0.038 | 0.070 | .996 |
| 28:7 | 2 | 4 | 0.120 | 0.069 | .462 |
| 28:7 | 3 | 4 | 0.083 | 0.032 | .107 |
| **Sham** | 1 | 2 | 0.036 | 0.070 | .997 |
| Sham | 1 | 3 | 0.116 | 0.091 | .780 |
| Sham | 1 | 4 | 0.158 | 0.089 | .451 |
| Sham | 2 | 3 | 0.080 | 0.070 | .851 |
| Sham | 2 | 4 | 0.123 | 0.069 | .443 |
| Sham | 3 | 4 | 0.043 | 0.032 | .730 |

**Table S4E: Multivariate** tests for the simple effect of Phase within each group in Figures 4 b and c.

| **Multivariate Tests** | | | | | | |
| --- | --- | --- | --- | --- | --- | --- |
| Group | | Value | F | Hypothesis df | Error df | Sig. |
| Anat | Pillai's trace | .089 | .453^a^ | 3.000 | 14.000 | .719 |
|  | Wilks' lam16:2a | .911 | .453^a^ | 3.000 | 14.000 | .719 |
|  | Hotelling's trace | .097 | .453^a^ | 3.000 | 14.000 | .719 |
|  | Roy's largest root | .097 | .453^a^ | 3.000 | 14.000 | .719 |
| 50:5 | Pillai's trace | .542 | 5.513^a^ | 3.000 | 14.000 | .010 |
|  | Wilks' lam16:2a | .458 | 5.513^a^ | 3.000 | 14.000 | .010 |
|  | Hotelling's trace | 1.181 | 5.513^a^ | 3.000 | 14.000 | .010 |
|  | Roy's largest root | 1.181 | 5.513^a^ | 3.000 | 14.000 | .010 |
| 28:7 | Pillai's trace | .357 | 2.594^a^ | 3.000 | 14.000 | .094 |
|  | Wilks' lam16:2a | .643 | 2.594^a^ | 3.000 | 14.000 | .094 |
|  | Hotelling's trace | .556 | 2.594^a^ | 3.000 | 14.000 | .094 |
|  | Roy's largest root | .556 | 2.594^a^ | 3.000 | 14.000 | .094 |
| Sham | Pillai's trace | .237 | 1.450^a^ | 3.000 | 14.000 | .271 |
|  | Wilks' lam16:2a | .763 | 1.450^a^ | 3.000 | 14.000 | .271 |
|  | Hotelling's trace | .311 | 1.450^a^ | 3.000 | 14.000 | .271 |
|  | Roy's largest root | .311 | 1.450^a^ | 3.000 | 14.000 | .271 |

**Table S5A:** Between-group comparisons of Normalized DA from Figures 5a and 5c.

| **(I) Group** | **(J) Group** | **Mean Difference (I–J)** | **Std. Error** | **Sig. (b)** |
| --- | --- | --- | --- | --- |
| BT | FBT | 0.014 | 0.046 | 1.000 |
| BT | FGT | –0.277* | 0.049 | < .001 |
| BT | GT | –0.532* | 0.046 | < .001 |
| FBT | FGT | –0.291* | 0.051 | < .001 |
| FBT | GT | –0.546* | 0.048 | < .001 |
| FGT | GT | –0.255* | 0.051 | < .001 |

**Table S5B:** Normalized DA across groups and phases in Figure 5a and 5c.

| **Estimates** | | | |
| --- | --- | --- | --- |
| Measure: Normalized DA | | | |
| Group | Phase | Mean | Std. Error |
| BT | 1 | 1.000 | .000 |
|  | 2 | .750 | .059 |
|  | 3 | .568 | .042 |
|  | 4 | .404 | .043 |
| FBT | 1 | 1.000 | .000 |
|  | 2 | .748 | .065 |
|  | 3 | .528 | .046 |
|  | 4 | .390 | .047 |
| FGT | 1 | 1.000 | .000 |
|  | 2 | 1.016 | .073 |
|  | 3 | .955 | .052 |
|  | 4 | .860 | .053 |
| GT | 1 | 1.000 | .000 |
|  | 2 | 1.302 | .065 |
|  | 3 | 1.284 | .046 |
|  | 4 | 1.265 | .047 |

**Table S5C:** Post hoc between-group comparisons of Normalized DA at each phase in Figure 5a and 5c.

| **Phase** | **(I) Group** | **(J) Group** | **Mean Difference (I–J)** | **Std. Error** | **Sig. (b)** |
| --- | --- | --- | --- | --- | --- |
| **1** | BT | FBT | –1.480E-16 | 0.000 | . |
|  | BT | FGT | –1.480E-16 | 0.000 | . |
|  | BT | GT | –3.257E-16 | 0.000 | . |
|  | FBT | FGT | 0.000 | 0.000 | . |
|  | FBT | GT | –1.776E-16 | 0.000 | . |
|  | FGT | GT | –1.776E-16 | 0.000 | . |
| **2** | BT | FBT | 0.002 | 0.088 | 1.000 |
|  | BT | FGT | –0.265 | 0.094 | 0.071 |
|  | BT | GT | –0.552* | 0.088 | < .001 |
|  | FBT | FGT | –0.268 | 0.098 | 0.083 |
|  | FBT | GT | –0.554* | 0.092 | < .001 |
|  | FGT | GT | –0.286 | 0.098 | 0.057 |
| **3** | BT | FBT | 0.040 | 0.062 | 0.989 |
|  | BT | FGT | –0.387* | 0.067 | < .001 |
|  | BT | GT | –0.716* | 0.062 | < .001 |
|  | FBT | FGT | –0.428* | 0.069 | < .001 |
|  | FBT | GT | –0.756* | 0.065 | < .001 |
|  | FGT | GT | –0.329* | 0.069 | 0.001 |
| **4** | BT | FBT | 0.015 | 0.064 | 1.000 |
|  | BT | FGT | –0.456* | 0.068 | < .001 |
|  | BT | GT | –0.861* | 0.064 | < .001 |
|  | FBT | FGT | –0.470* | 0.071 | < .001 |
|  | FBT | GT | –0.875* | 0.067 | < .001 |
|  | FGT | GT | –0.405* | 0.071 | < .001 |

**Table S5D:** Post hoc within-group comparisons of Normalized DA across phases from Figure 5a and c

| **Group** | **(I) Phase** | **(J) Phase** | **Mean Difference (I–J)** | **Std. Error** | **Sig. (b)** |
| --- | --- | --- | --- | --- | --- |
| **BT** | 1 | 2 | 0.250* | 0.059 | 0.004 |
|  | 1 | 3 | 0.432* | 0.042 | < 0.001 |
|  | 1 | 4 | 0.596* | 0.043 | < 0.001 |
|  | 2 | 3 | 0.182* | 0.045 | 0.006 |
|  | 2 | 4 | 0.346* | 0.053 | < 0.001 |
|  | 3 | 4 | 0.164* | 0.034 | 0.001 |
| **FBT** | 1 | 2 | 0.252* | 0.065 | 0.008 |
|  | 1 | 3 | 0.472* | 0.046 | < 0.001 |
|  | 1 | 4 | 0.610* | 0.047 | < 0.001 |
|  | 2 | 3 | 0.220* | 0.050 | 0.003 |
|  | 2 | 4 | 0.358* | 0.058 | < 0.001 |
|  | 3 | 4 | 0.138* | 0.037 | 0.012 |
| **FGT** | 1 | 2 | –0.016 | 0.073 | 1.000 |
|  | 1 | 3 | 0.045 | 0.052 | 0.953 |
|  | 1 | 4 | 0.140 | 0.053 | 0.103 |
|  | 2 | 3 | 0.060 | 0.056 | 0.877 |
|  | 2 | 4 | 0.155 | 0.065 | 0.166 |
|  | 3 | 4 | 0.095 | 0.042 | 0.203 |
| **GT** | 1 | 2 | –0.302* | 0.065 | 0.002 |
|  | 1 | 3 | –0.284* | 0.046 | < 0.001 |
|  | 1 | 4 | –0.265* | 0.047 | < 0.001 |
|  | 2 | 3 | 0.018 | 0.050 | 1.000 |
|  | 2 | 4 | 0.037 | 0.058 | 0.990 |
|  | 3 | 4 | 0.019 | 0.037 | 0.997 |

**Table S5E: Multivariate** tests for the simple effect of Phase within each group in Figures 5a and 5c.

| **Multivariate Tests** | | | | | | |
| --- | --- | --- | --- | --- | --- | --- |
| Group | | Value | F | Hypothesis df | Error df | Sig. |
| BT | Pillai's trace | .929 | 60.858^a^ | 3.000 | 14.000 | <.001 |
|  | Wilks' lambda | .071 | 60.858^a^ | 3.000 | 14.000 | <.001 |
|  | Hotelling's trace | 13.041 | 60.858^a^ | 3.000 | 14.000 | <.001 |
|  | Roy's largest root | 13.041 | 60.858^a^ | 3.000 | 14.000 | <.001 |
| FBT | Pillai's trace | .922 | 55.404^a^ | 3.000 | 14.000 | <.001 |
|  | Wilks' lambda | .078 | 55.404^a^ | 3.000 | 14.000 | <.001 |
|  | Hotelling's trace | 11.872 | 55.404^a^ | 3.000 | 14.000 | <.001 |
|  | Roy's largest root | 11.872 | 55.404^a^ | 3.000 | 14.000 | <.001 |
| FGT | Pillai's trace | .392 | 3.007^a^ | 3.000 | 14.000 | .066 |
|  | Wilks' lambda | .608 | 3.007^a^ | 3.000 | 14.000 | .066 |
|  | Hotelling's trace | .644 | 3.007^a^ | 3.000 | 14.000 | .066 |
|  | Roy's largest root | .644 | 3.007^a^ | 3.000 | 14.000 | .066 |
| GT | Pillai's trace | .724 | 12.272^a^ | 3.000 | 14.000 | <.001 |
|  | Wilks' lambda | .276 | 12.272^a^ | 3.000 | 14.000 | <.001 |
|  | Hotelling's trace | 2.630 | 12.272^a^ | 3.000 | 14.000 | <.001 |
|  | Roy's largest root | 2.630 | 12.272^a^ | 3.000 | 14.000 | <.001 |
